# An Independent Coding Scheme for Distance versus Position in the Hippocampus

**DOI:** 10.1101/2024.12.27.629676

**Authors:** Mathilde Nordlund, Nicolas Levernier, Massimiliano Trippa, Romain Bourboulou, Geoffrey Marti, Rémi Monasson, Hervé Rouault, Jérôme Epsztein, Julie Koenig-Gambini

## Abstract

Animals navigate using cognitive maps of their environment, integrating external landmarks and distances between them. Hippocampal place cells are the neuronal substrate of these cognitive maps. However, while hippocampal allocentric position coding in reference to external landmarks is well characterized, the determinants of idiothetic hippocampal distance coding remain poorly understood. Using virtual reality, electrophysiological recordings in mice, and local cue manipulations we could dissociate distance from position coding. In the cue-poor condition, we found pervasive distance coding with high distance indices in all bidirectional place cells including both superficial and deep CA1 pyramidal cells. In this condition, the mapping of distance onto a low-dimensional manifold and rigid distance relationships between place fields suggested strong attractor dynamics similar to those observed for grid cells. Inactivation of the medial septum (MS), which disrupts grid cells, significantly reduced both distance coding and rigid distance dynamics, suggesting an alteration (but not complete abolition) of the underlying attractor. In contrast, allocentric position coding could be observed in cue-rich environments, predominantly engaged deep CA1 pyramidal cells, and persisted during MS inactivation. These results are consistent with a selective contribution of grid cells and associated rigid attractor dynamics to hippocampal idiothetic distance coding but not allocentric position coding.

## INTRODUCTION

When navigating to find food, escape predators, or find potential mates, animals can use two strategies: the cognitive map and path integration^1^. The cognitive map strategy relies on the acquisition, through learning, of an internal representation of the environment that contains spatial relationships between significant landmarks and important locations. Animals can locate themselves on this map using a variety of external cues (allocentric position coding). The hippocampus has been implicated in this type of navigation^2^. During navigation, hippocampal principal cells, called place cells, increase their activity each time an animal is in the same location in reference to external cues^3^. Accordingly, the study of allocentric position coding have dominated the field of hippocampal research for several decades^4^. In contrast, the path integration strategy allows animals to locate themselves independently from external sensory cues using self-motion cues to continuously update their location along a travel path by integrating the distance traveled in different direction^5^. However, while decades of research have revealed how the brain represents absolute location (i.e. hippocampal position coding), reports of hippocampal distance coding in the literature have been scarce^6–8^. Accordingly, the neural mechanisms of hippocampal distance coding and whether they differ from position coding remain largely unknown.

At the cellular level, while the morpho-functional heterogeneity of CA1 pyramidal cells along the hippocampal radial axis is now well established^9–11^, it remains unknown whether distance coding can be mapped specifically to a given CA1 pyramidal cell subtype (deep versus superficial). At the network level, grid cells were initially proposed to contribute to hippocampal position coding. However grid fields show fixed and rigid distance relationship^12,13^, supported by a recurrent continuous attractor network with a toroidal topology^14^, and could provide a distance metric to place cells^15^. In addition, medial septum (MS) inactivation, which reduces theta rhythmicity and selectively alters the spatial firing pattern of grid cells^16,17^ does not prevent place cells’ position coding^17,18^ but disrupts distance estimation based on path integration in rats navigating in the dark^19^ and place cell activity away from external landmarks^18,20^. Finally, during postnatal development, the accuracy of place cell coding away from boundaries improves with the functional maturation of grid cells^21^. Taken together, these results are consistent with a specific contribution of grid cells to hippocampal distance coding^22^. However, grid cell firing is disrupted in the absence of visual cues when self-motion-based distance coding should dominate^23,24^, and distance coding can be observed in preweaned rats before grid cell maturation^25^, calling this specific role into question.

To address these questions, we performed large-scale extracellular recordings from CA1 pyramidal cells during virtual navigation in cue-rich and/or cue-poor virtual environments^26^. We focused on bidirectional place cells in hippocampal area CA1 with place fields in both forward and backward trials to decipher whether individual cells exhibited position coding (firing at the same position with respect to external visual cues in forward and backward trials) or distance coding (firing at the same distance from the start of each trial)^6,27^. Distance coding dominated hippocampal activity in the absence (cue-poor), but not in the presence (cue-rich), of salient local visual cues (virtual 3D objects). This ability to isolate hippocampal distance coding in the cue-poor condition provides a unique opportunity to decipher its cellular and circuit determinants. In particular, we asked whether it is supported by a specific subpopulation of deep or superficial CA1 pyramidal cells. We also characterized distance coding at the population level, looking for signatures of attractor dynamics using dimensionality reduction approaches and correlation analysis. Finally, we tested the effect of MS inactivation on hippocampal distance coding at both the cellular and population level. Overall, our results demonstrate pervasive egocentric distance coding in the hippocampus during virtual navigation in cue-poor environments, supported by both deep and superficial CA1 pyramidal cells. In this condition, place cell distance coding is strongly reduced (but not completely abolished) by MS inactivation. However, position coding was still observed in a cue-rich environment under MS inactivation. Taken together, our results are consistent with a preferential contribution of grid cells to hippocampal distance but not position coding.

## RESULTS

### Absence of local visual cues promotes distance coding in bidirectional CA1 place cells

Proximal visual cues have a strong influence on the coding of hippocampal place cells in virtual reality environments^26^. We first tested their influence on the propensity of hippocampal place cells to perform position or distance coding. For this, head-fixed mice were trained to run in 2 m long virtual linear tracks locally enriched with visual cues (two or three virtual 3D objects; cue-rich track) or without objects (cue-poor track; Figures 1A and 1B) for sweetened water rewards distributed at the track ends. Immediately after reward consumption, mice were teleported in the same position but facing the opposite direction of the track allowing them to run back and forth in the same environment. Once animals reached a stable and proficient behavior (at least one reward/min during a 60-min session: 4.38 ± 0.52 cue-rich and 5.17 ± 0.61 cue-poor, t_26_ = 0.82, p = 0.42, unpaired t-test), we recorded spiking activity in the pyramidal cell layer of the CA1 hippocampal region using 8-shanks silicon probes in the right and/or left hemisphere over the course of 2–3 days (cue-rich: 402 neurons from 9 recording sessions in 4 mice; cue-poor: 1462 neurons from 19 recording sessions in 14 mice).

**Figure 1:**
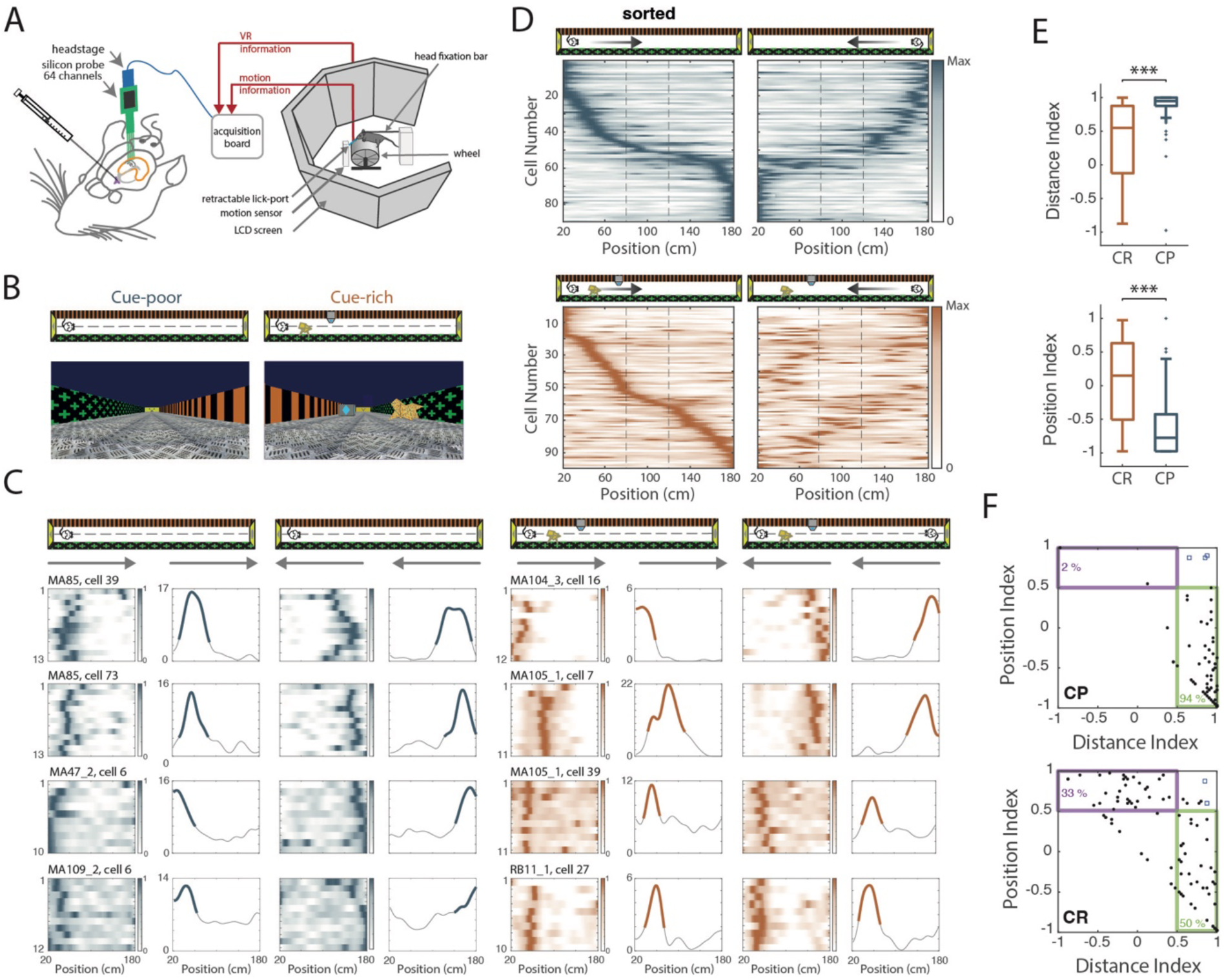
Absence of local visual objects promotes distance coding in CA1 bidirectional place cells. (**A**) Schematic of the virtual reality set up. The mouse is head-fixed and placed on a wheel surrounded by LCD screens displaying a virtual environment. A guide cannula is implanted in the medial septum for intracranial infusion of muscimol or aCSF (see Figure 3). (**B**) Top and first-person views of the virtual linear tracks used. Left: cue-poor track (CP) and right: cue-rich track with 2 virtual 3D objects (CR). (**C**) 4 examples of bidirectional cells recorded in the cue-poor track (left) or in the cue-rich track (right). For each cell, the panel shows the color-coded firing rate map for successive trials in both directions (arrows) as a function of the position in the maze and the corresponding mean firing rate by position (reward zones excluded). Bold traces indicate positions of the detected place field. (**D**) Color-coded mean firing rate maps of place fields detected in the forth trials (left, sorted by the location of place fields peak) and back trials (right, sorted according to firing order in the forth trials) of all bidirectional cells recorded in the track with objects (top) or without objects (bottom). The color represents for the intensity of the mean firing rate in a given bin normalized to the maximum mean firing rate (peak rate) in each direction. Note that in the cue-poor condition, the mean firing map in one direction is the mirror image of the mean firing map in the other direction, indicating strong distance coding. Dotted lines indicate the central zone of the maze. Bidirectional cells with both fields located in this region were excluded from the analysis. (**E**) Box plots comparing distance (top) or position (bottom) indices between CR and CP. (**F**) Scatter plots of distance index versus position index for each bidirectional cell recorded in CP (top) or CR (bottom). For each track condition, cells exhibiting position or distance coding are indicated by a purple or green box, respectively. The plots also show the percentage of bidirectional cells with distance or position coding. Note that in these plots, by definition, no point can be observed below the top-left-bottom-right diagonal.

We focused on bidirectional place cells because these cells, that have place fields in back-and-forth trials, allows distinguishing position coding (firing at the same position with respect to external visual cues in forward and backward trials) from distance coding (firing at the same distance from the start of each trial)^6,27^.

The percentage of bidirectional place cells was higher in the presence of proximal sensory cues (46.1 ± 4.7 % in the cue-rich versus 30.5 ± 4.0 % in the cue-poor track; t_26_ = −2.33, p = 0.028, two-tailed unpaired t test, Figure S1C)^27^. In the cue-poor environment, the vast majority of bidirectional place cells appeared to have place fields at a similar distance from the start of each journey in back-and-forth trials (Figure 1C, blue). This was evident when we plotted the place fields sorted by their location in one direction (Figure 1D, “sorted”) and compared them with the location of the place fields when mice ran in the opposite direction (Figure 1D): we observed a mirror image. To quantify this, we computed a distance index (see methods) whose value vary from 1 when place fields are at the exact same distance in back-and-forth trials to −1 when they are at the same position. In the cue-poor track, the distance index was close to 1 (0.88 ± 0.03; n = 86 bidirectional cells; Figures 1E).

Next, we focused on bidirectional place cells in the cue-rich track. As previously reported, the presence of proximal visual cues significantly increased the proportion of spatially modulated cells (place cells) among active cells (cue-rich: 68.9 ± 4.6 % vs cue-poor: 32.2 ± 3.1 %; t_26_ = −6.42 p < 10^-6^, two-tailed non-paired t test, Figure S1A, B)^26,28^. In this condition we also observed bidirectional distance coding place cells (Figure 1C, orange and 1D), but in addition, some bidirectional place cells also appeared to code at the same position with respect to external cues in back-and-forth trials (Figure 1C, orange and 1D). Indeed, the distance index was significantly lower in the cue-rich compared to the cue-poor environment (0.38 ± 0.05; Z = 7.08, p<10^11^, two-tailed Wilcoxon Rank Sum test or WRS test; Figure 1E) while the position index was significantly higher in the in cue-rich compared to the cue-poor environment (CR: 0.06 ± 0.07; CP: −0.62 ± 0.05; Z = −6.66, p<10^10^, two-tailed WRS test; Figure 1E).

Next, we classified bidirectional place cells as distance-coding cells or position-coding cells (see Methods). Strikingly, in the cue-poor track almost all bidirectional place cells were distance-coding cells: 94.2 % (n = 81 out of 86) with only 2.3 % (n = 2) of bidirectional place cells showing position coding (Figures 1D and 1F; Supplementary Figure 1D). In the cue-rich track, the proportion of distance-coding and position-coding cells was more balanced: Distance-coding: 50.5 % (n = 49 out of 97); position-coding: 33.0 % (n = 32) (Figures 1D and 1F; Supplementary Figure 1E). The proportions of cells coding for position and distance were significantly different between track conditions (X_2_ (2, n = 183) = 42.7, p<10^9^, chi-squared test).

We concluded that distance coding dominates bidirectional place cell activity in virtual reality environments in the absence of local visual cues but that a mixture of position and distance coding is observed in heterogeneous virtual environments enriched with local visual cues.

Objects are heterogeneously distributed in the cue-rich track with one part enriched with objects (cue-rich part) and another part without objects (cue-poor part). Since both distance and position coding place cells can be recorded in this track, we wondered whether their place fields were heterogeneously distributed between cue-rich and cue-poor parts of the track. Interestingly the place fields of distance-coding cells were equally distributed between cue-rich and cue-poor parts of the track (18.4 % cue-rich vs 19.4 % cue-poor), whereas the place fields of position-coding cells were preferentially located in the cue-rich part of the track (59,4 % vs 14.1 % cue-poor; X_2_ (1, n = 146) = 15.14, p<10^4^, Chi-square test, Figure S1F).

Overall, these results show that in the cue-rich track, place cells can exhibit distance coding in both cue-rich and cue-poor parts of the track, whereas position coding is mostly observed in cue-rich parts of the track.

### Both deep and superficial CA1 pyramidal cells exhibit distance coding in cue-poor environments

CA1 pyramidal cells are morpho-functionally diverse depending on their location along radial, proximo-distal and transverse axes of the hippocampus^29–32^. In the radial axis, CA1 pyramidal cells whose soma is located deep in the layer (closer to the stratum oriens) are more strongly activated by local sensory cues and have their place fields in cue-rich locations in heterogeneous environments. Conversely, CA1 pyramidal cells whose soma is located more superficially in the layer (closer to stratum radiatum) are less activated by local sensory cues and have their place fields enriched in cue-poor locations^11,20,32^. Next, we asked how hippocampal distance/position coding map onto this heterogeneity.

To decipher the anatomical distribution of cells along the CA1 radial axis, we defined the middle of the CA1 pyramidal layer as the recording site where sharp-wave ripples had the largest amplitude ^29,32^. Cells above or below this threshold were classified as deep or superficial, respectively (Figure S2A). We were able to record a large number of deep and superficial pyramidal cells when mice explored both cue-poor and cue-rich tracks (cue-poor: 362 deep and 642 superficial cells, 14 sessions in 8 mice; cue-rich: 122 deep and 246 superficial cells, 8 sessions in 3 mice). Overall, there was no difference in the propensity of deep and superficial cells to be active and spatially modulated in either cue-poor or cue-rich environments (Figure S2B and S2C).

We then compared distance/position coding of deep and superficial CA1 pyramidal cells in cue-poor and cue-rich environments focusing on bidirectional place cells (Figures 2A and 2B). The percentage of bidirectional cells among all place cells did not differ between deep and superficial pyramidal cells (Figure S2D). In the cue-poor environment the majority of both deep and superficial cells had place fields at the same distance from the start of each journey (i.e. distance-coding cells; deep: 87.5 %; superficial: 90.7 %, X^2^(2, n = 67) = 1.83, p=0.40, Chi-square test). Accordingly, distance coding indices were similarly high for both cell types (deep: 0.79 ± 0.09; superficial: 0.91 ± 0.02, respectively; Z = −1.45, p = 0.15, two-tailed Wilcoxon Rank Sum test, Figure 2C). Thus, in the absence of local sensory cues distance coding dominates in both deep and superficial bidirectional CA1 pyramidal cells.

**Figure 2:**
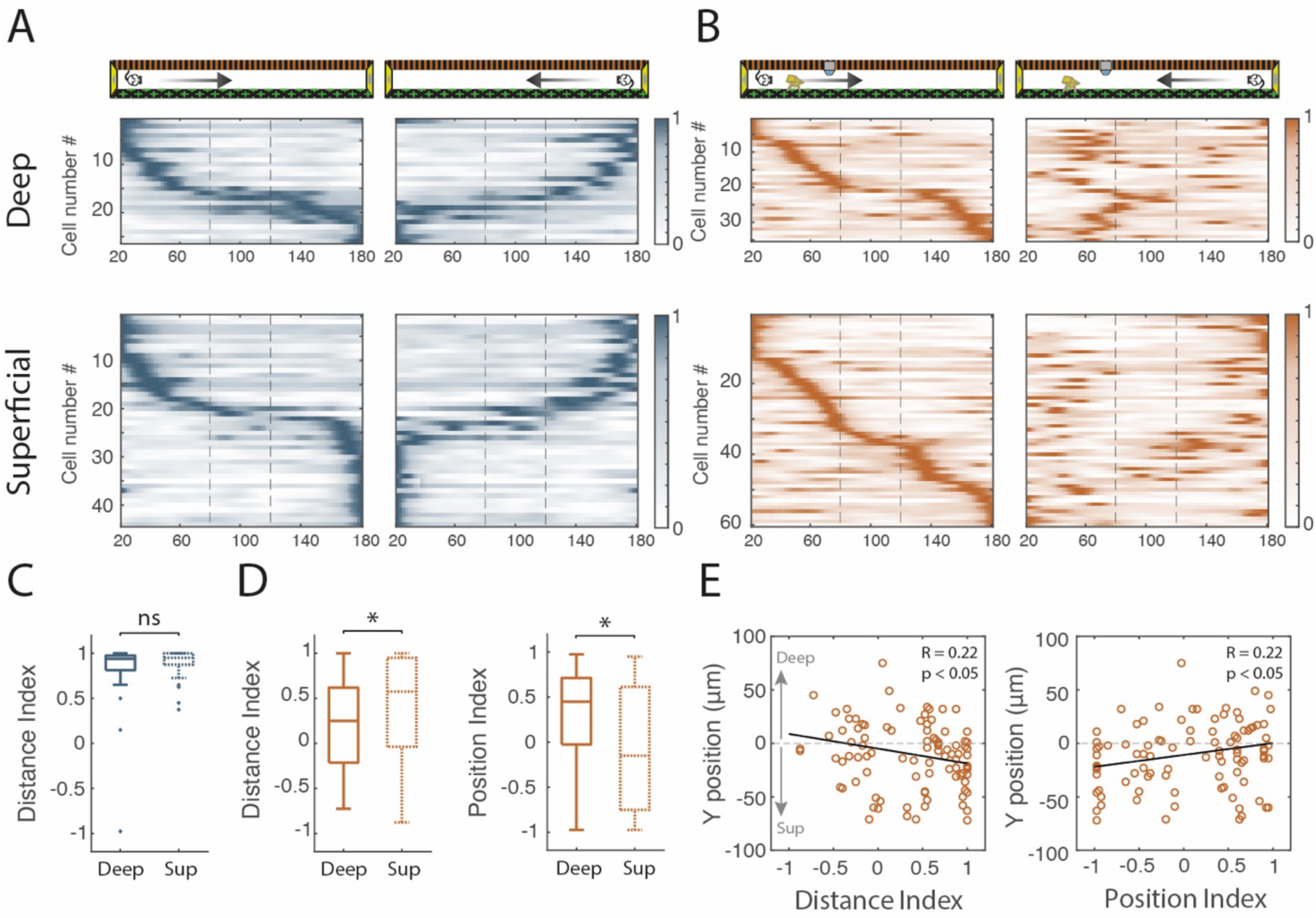
Distance and position coding in deep and superficial pyramidal cells. (**A**) Color-coded mean firing rate maps in the forth (left) and back (right) trials of deep (top) and superficial (bottom) bidirectional place cells recorded in the cue-poor track. The color code represents the normalized mean firing rate in a given bin. Place cells are ordered according to the position of their peak firing rate in the forth direction (reward zones excluded). (**B**) Color-coded mean firing rate maps in the forth (left) and back (right) trials of deep (top) and superficial (bottom) bidirectional place cells recorded in the cue-rich track. (**C**) Box plots comparing the distance indices between deep and superficial cells in the cue-poor track. (**D**) Box plots comparing distance index (left) or position index (right) between deep and superficial cells in the cue-rich track. (**E**) Scatter plots of distance index (left) or position index (right) in the cue-rich track versus Y-position in the cell layer (μm).

In the heterogeneous cue-rich environment, however, the distance index of superficial cells was significantly higher than that of deep cells (superficial: 0.46 ±0.07 and deep: 0.23 ±0.08, respectively, Z = −2.2251, p = 0,0261, two-tailed WRS test, Figure 2D). Furthermore, we observed a significant negative correlation between cell body depth and distance index (R = −0.22, p < 0.05, Pearson correlation, Figure 2E). Accordingly, in this track, the position index was higher for deep cells (deep: 0.28 ±0.09; superficial: −0.05 ± 0.09, Z = 2.0937, p = 0.0363, two-tailed WRS test, Figure 2D) and the correlation between cell body depth and position index was significantly positive (R = 0.22 p < 0.05, Pearson correlation, Figure 2E). Nevertheless, both deep and superficial pyramidal cells could be classified as either distance-coding cells or position-coding cells. When considering the superficial cell population, the proportion of distance-coding cells (55.0%) tended to be higher than the proportion of position-coding cells (28.3%) but without reaching statistical significance. On the other hand, when considering the deep cell population, the proportion of distance coding (42.9 %) and position coding (42.9 %) cell was identical (X^2^(2, n = 95) = 2.11, p = 0.35, chi-squared test).

Taken together these results show that, in the absence of local visual cues, both CA1sup and CA1deep are equally recruited to code distance. However, in a heterogeneous track locally enriched with visual cues, our results showed a bias towards distance coding in CA1sup cells and towards position coding in CA1deep cells.

### Distance coding in unidirectional place cells and non-place cells

Distance coding dominates the activity of bidirectional place cells in the cue-poor condition. What about unidirectional place cells which have a place field when animals navigate in one but not the opposite direction^6,27^ and represent the majority of spatially modulated cells in linear tracks? In our recordings, unidirectional place cells fired at similar rates in both directions (firing rate: Z = 0.93, p = 0.086, two-tailed WSR test, Figure S3A; spatial information: Z = 0.90, p = 0.37, two-tailed WSR test, Figure S3A), but less reliably (stability: Z = 4.04, p < 10^4^, two-tailed WSR test; Figure S3A). We therefore looked for traces of distance coding in these cells. Looking at the raster plots of individual cells an increased firing rate was often observed in the opposite direction at the same distance as the place field from the start of the journey, reminiscent of distance coding (Figure S3B). This was also evident when all place fields were plotted ordered by their location in the coding direction (Figure S3C, left sorted) and compared with locations associated with increased activity (dark blue) in the opposite direction (Figure S3C). To quantify this, we calculated the spatial correlation between the mean firing rate vectors in the two directions aligned to the start of each journey. The spatial correlation of unidirectional cells reached a mean value of 0.57 ± 0.02, significantly higher than a shuffled dataset (where pairs of cells in back-and-forth trials were randomized see Method; Figure S3D) but lower than the spatial correlation in bidirectional cells (0.84 ± 0.02; X^2^(3) = 505.5, p < 10^−108^; Kruskal Wallis, p < 10^5^ in both cases, multiple comparisons). Non-place cells (with no place fields in any direction) also showed a spatial correlation significantly higher than shuffle (0.29 ± 0.01 vs −0.023 ± 0.01, p < 10^8^, multiple comparison), suggesting that some distance coding could also be detected within this population.

We conclude that in cue-poor environments, distance coding can also be detected in unidirectional and non-place cells suggesting a global coding at the population level.

### Population analysis of hippocampal distance coding

Our previous results show that in the absence of local visual cues, distance coding dominates hippocampal place cells activity. Hippocampal distance coding is readily observable in bidirectional place cells, with place fields at the same distance from the start of each journey, but also in unidirectional place cells and non-place cells, suggesting a predominant distance coding at the population level. To better visualize the population dynamics in the cue-poor track we applied nonlinear dimensionality reduction technique (UMAP, see Methods) to the spiking activity of all recorded active cells for each recording session in this condition (Figure 3A). Projection onto a 2D plane revealed a cloud of points with an elongated shape (1D manifold), each point corresponding to a population vector summarizing hippocampal network activity at a given time (Figure 3A). Distance from the start of each journey, but not position, nicely mapped onto the coordinate of this manifold for a majority of recording sessions in this condition (Figure 3A), confirming the predominance of distance coding at the population level in the cue-poor track. Several CA1 pyramidal cells showed consistently increased activity at restricted locations on the manifold reminiscent of the distribution of place cell activity along the cue-poor track (Figure 3B). Indeed, there was a good correlation between the distance traveled by the animal from the start of each journey and the evolution of the population activity along the manifold (Figure 3C). The reduction of hippocampal activity onto a low-dimensional manifold is consistent with continuous attractor dynamics. These dynamics could impose internal constraints on the neuronal activity of place cells such as rigid distance relationships between place fields as recently described for grid cells^12,14,33^. To test whether similar dynamics might underlie relative distance coding, we looked for rigid distance relationships between hippocampal firing across laps in the cue-poor condition. Despite the large intertrial spatial variability of place fields in this condition^26^, spatial cross-correlation analysis revealed that a significant fraction of cell pairs maintained a fixed distance between their place fields across laps (Figure 3D, E). The correlation matrix showed that short distances between place fields (in the range of 10-20 cm) were overrepresented (Figure 3F). A 2D multidimensional scaling on the distance matrix showed that spatial shifts formed a mostly linear representation compatible with an underlying 1D manifold (Figure 3G), consistent with the UMAP analysis.

**Figure 3:**
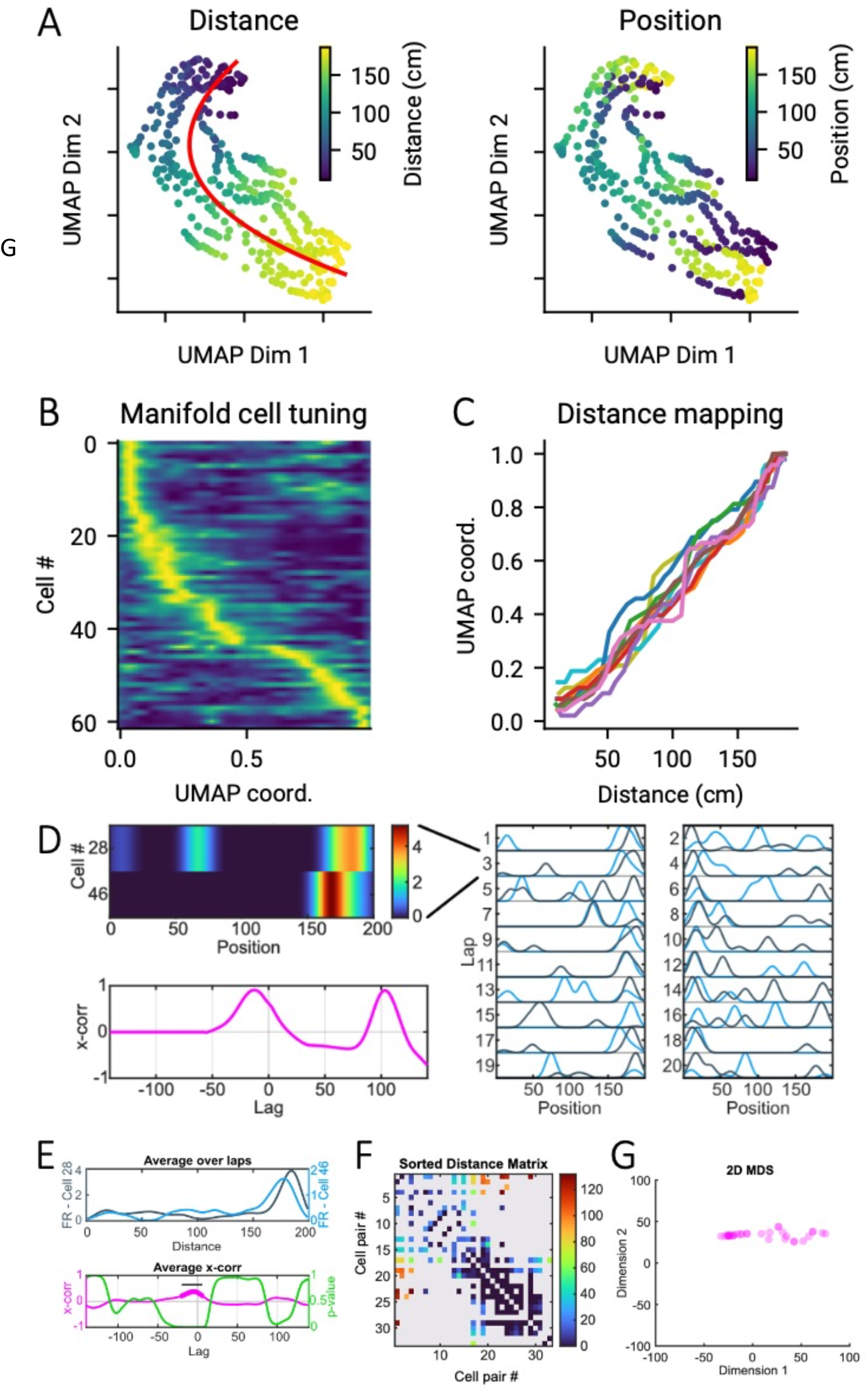
Attractor dynamics of hippocampal distance coding. (**A-B**) Example of UMAP projection of hippocampal activity during a recording session in the cue-poor track, colored according to the distance with respect to the start of the lap (**left**) or the position (right). (**B**) Example of localized increased in activity sorted according to UMAP coordinate. (**C**) Correlation between distance from the start of the maze and UMAP coordinate (UMAP Coord.) for several laps during the recording session in A. (**D**) Left: Firing rate (top, color coded) and firing cross correlation (bottom) between cell #28 and cell #46 during a single forward lap in the cue-poor maze. Right: Normalized average firing of cell #28 (black) and cell #46 (blue) for all laps in the forward (odd) and backward (even) directions. (**E**) Average firing rate of cell #28 (black) and cell #46 (blue) in the maze plotted against distance from the start (top) and corresponding cross-correlation (pink) and associated p values (green) for each distance lag (bottom). The thicker pink line identifies ranges of distance lags with p-values lower than 0.001. The black bar locates a cluster of at least 10 significant adjacent lags. (**F**) Pairs of cells with significant cross-correlations for an example mouse; only the cells involved in at least one selected pair are included. Color coding represents the distance lag (in cm) encoded by each cell pair. (**G**) 2D multidimensional scaling dimensionality reduction applied to the distance matrix in panel F. Note the linear trend consistent with the one-dimensional nature of the environment.

Population level analysis therefore confirms that distance coding dominates hippocampal activity at the population level and could result from attractor dynamics imposing rigid distance relationships.

### Medial septum inactivation alters hippocampal distance coding

Our previous results show that distance coding dominates CA1 pyramidal cells’ activity in the cue-poor track. At the population level, the mapping of distance coding onto a 1D dimensional manifold is compatible with attractor dynamics. Accordingly, we observed that the activity of a majority of cells is tuned to the 1D manifold coordinate and shows rigid distance relationship across trials. These properties are reminiscent of grid cell activity which also lies on a low-dimensional manifold that imposes rigid distance relationships between grid fields of different cells.

The spatial organization of grid cell firing fields, which is thought to support path integration, is altered by medial septum (MS) inactivation^16,17^. We next tested the impact of MS inactivation on hippocampal distance coding in the cue-poor environment. Hippocampal neuronal activity was first recorded while mice performed back-and-forth trials in a familiar cue-poor environment (at least 20 trials). We then inactivated the MS, by micro-infusion of muscimol, an agonist of GABA_A_ receptors (see Methods). Muscimol infusion drastically reduced the power of theta oscillations recorded in the hippocampus (by 83.7 ± 4.1 %; t_17_ = −11.21, p < 10^8^ vs infusion of artifical cerebrospinal fluid or aCSF, paired t-test, Figure S4A-C).

The performance of the mice in the task was slightly but significantly reduced under MS inactivation (rewards/min: 3.59 ± 0.72 before and 3.28 ± 0.72 after, t_6_ = 2.65, p = 0.038, paired t-test). There was no significant change in the mean velocity before and after muscimol (t_6_ = −0.25, p = 0.81, Figure S4D) but animals tended to make more stops per trial after the infusion (t_6_ = −2,32, p = 0.060, paired t-test; Figure S4E). However, this was likely due to the length of the entire procedure (see Methods) rather than the injection of muscimol per se, as a similar increase was observed after injection of aCSF (t_11_ = −3,73, p < 0.01, paired t-test; Figure S4E). The total number of pyramidal cells recorded before and during the MS inactivation was 495 and 500, respectively (n = 7 recording sessions in 5 mice). As expected, (Brandon et al., 2011; Koenig et al., 2011), muscimol infusion in the MS significantly decreased the firing rate of active cells (Z = 3.80, p < 10^3^, two-tailed WRS for active place cells; Z = 2.99, p = 0.003, two-tailed WRS for active non-place cells, Figure S5A). However, the proportion of active pyramidal cells was not different before and after muscimol (67.5 ± 5.1 % and 61.4 ± 1.8 %, respectively, Z = 1.25, p = 0.26, n = 7 sessions, two-tailed WRS, S5B). The proportion of place cells among active cells was also similar (32.4 ± 4.4 % and 21.2 ± 2.9 %, respectively, t6 = 1.93, p = 0.10, n = 7 sessions, paired t-test, Figure S5B). Spatial information (Figure S5C) was mostly preserved under muscimol with no significant change detected (0.26 ± 0.03 bit/spike before and 0.25 ± 0.04 bit/spike after, Z = −0.0012, p = 0.99; n = 146 place cells before and 65 after, two-tailed WRS) as well as out/in field firing ratio (0.50 ± 0.02 before and 0.53 ± 0.03 after; Z = −1.28, p = 0.20, two-tailed WRS). However, place cells were less stable after MS inactivation (0.24 ± 0.01 before and 0.13 ± 0.01 after, Z = 5.80, p < 10^8^, two-tailed WRS, Figure S5C). Altogether, we concluded that cells are less active and stable under MS inactivation even if no difference was observed in terms of spatial information.

We then asked whether MS inactivation could alter hippocampal distance coding in the cue-poor track. We indeed observed a drastic decrease in the number of distance-coding bidirectional place cells under MS inactivation (from 38 before to 5 after MS inactivation; 86.8 % decrease, Figure 4A). This loss of bidirectional distance coding place cells under MS inactivation also reached significance on a session-by-session analysis (t_6_ = 4.75, p = 0.0032, two-tailed paired t-test, Figure 4B). In comparison, the total number of unidirectional cells decreased less (by 21.4 %; 70 to 55 during the MS inactivation). We next compared the percentages of bidirectional and unidirectional cells, calculated from the total number of place cells, before and after the muscimol infusion. Before MS inactivation, 35.2 % of place cells were bidirectional and 64.8% were unidirectional, whereas after MS inactivation, only 8.3% of place cells were bidirectional and the remaining 91.7% were unidirectional (X2(1, n = 168) = 14.6, p = < 10^3^, chi-squared test; Figure 4B). To further estimate the effects of MS inactivation on bidirectional and unidirectional place cells, we calculated their relative change (N_after_-N_before_)/N_before_. For bidirectional place cells, the relative change was close to −1 and significantly different from 0, indicating a strong decrease during MS inactivation (mean: −0.88 ± 0.06, p = 0.016, n = 7, two-tailed WSR, Figure 4C). The relative change was however not significantly different from 0 for unidirectional cells (mean: −0.09 ± 0.13, t_6_ = −0.69, p = 0.51, one-sample t-test, Figure 4C) and significantly different than bidirectional cells (p = 0.0156, n = 7, two-tailed WSR, Figure 4C). Since most of the active cells are non-place cells in the cue-poor track, we also examined the effects on MS inactivation on this category of cells. The relative change for non-place cells was not significantly different from 0 (mean: 0.18 ± 0.19, t_6_ = 0.95, p = 0.38, one sample t-test, Figure 4D). Taken together, our results indicate that MS inactivation preferentially silenced distance-coding bidirectional place cells.

**Figure 4:**
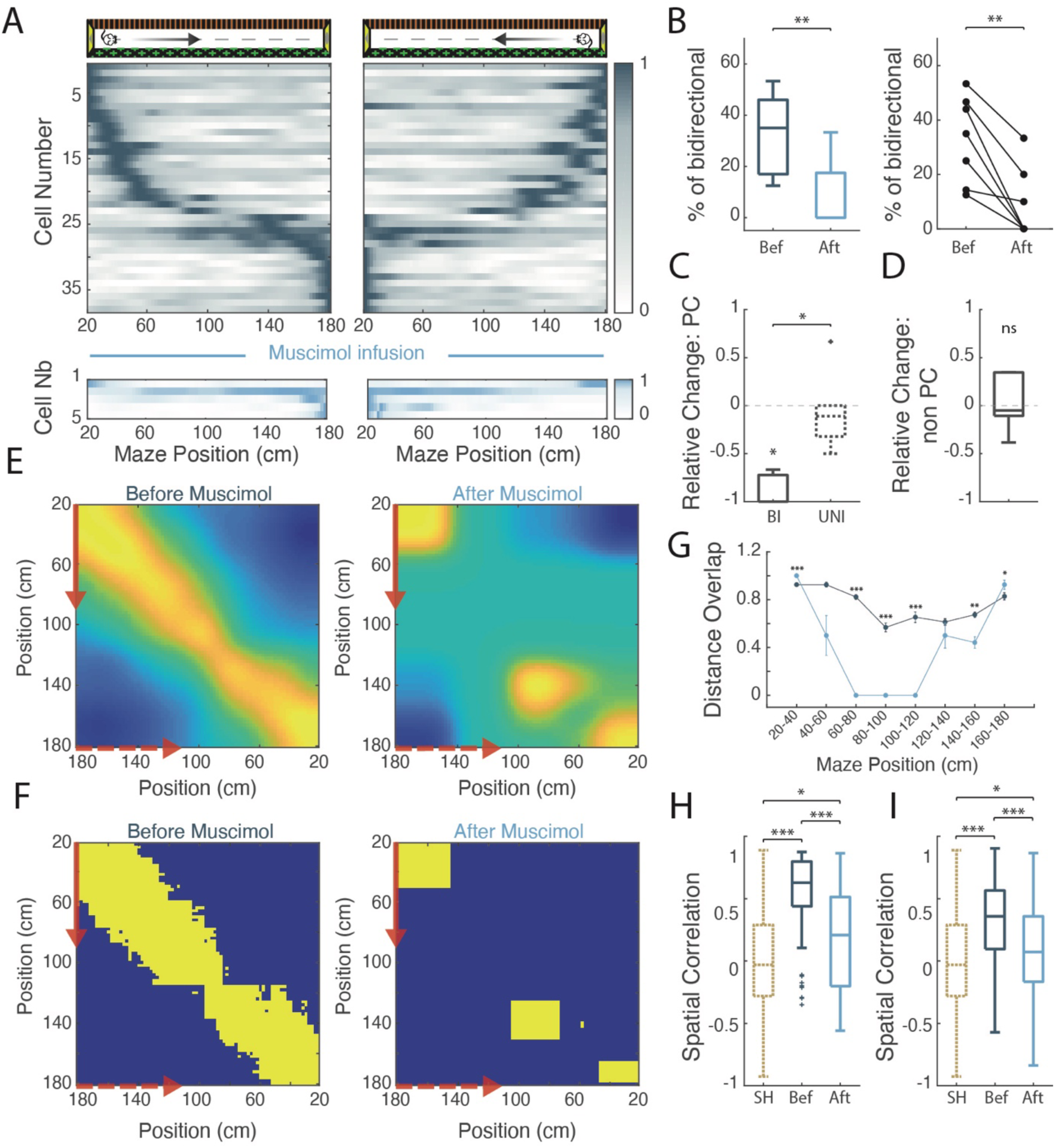
Hippocampal distance coding is altered under medial septum inactivation. (**A**) Color-coded mean firing rate maps in the forth (left) and back (right) trials of bidirectional cells recorded before (top, dark blue) and after (bottom, light blue) the muscimol infusion in the cue-poor track. The color codes represent the intensity of the mean firing rate of the place cell, normalized to the maximum mean firing rate (peak rate) in each direction. Place cells in both directions are ordered according to the position of their peak rate in the forth trials (reward zones excluded). (**B**) Left: Box plot of the percentage of bidirectional place cells recorded in the cue-poor track before and during the MS inactivation. Right: Mean (±SEM) percentage of bidirectional cells before and during MS inactivation for each recording session with at least 4 place cells. (**C**) Box plots of relative change (= number of cells after minus number of cells before, divided by number of cells before) of bidirectional (left) and unidirectional (right) place cells. The asterisk indicates that the relative change for the bidirectional cells was significantly different from 0 (p<0.05). (**D**) Box plot of relative change for active non-place cells. (**E**) Population vector correlations, at each position bin, between place field presence in bidirectional cells recorded in the cue-poor track (reward zones excluded) in both directions before (left) and during (right) MS inactivation. For each axis, the arrows indicate the trial direction (solid red for forward trials and dashed red for backward trials). The intensity of the correlation is color-coded from highest (yellow) to lowest (blue). Prior to MS inactivation, the presence of a place field was highly correlated with the presence of a place field at equivalent locations in the opposite direction, oriented to the starting point. During MS inactivation, high correlations were lost in the middle part of the trace. (**F**) Population vector matrices showing bins of correlation above the 99^th^ percentile threshold of the shuffled data before (left) and during (right) MS inactivation. (**G**) Distance overlap of bidirectional place cells in 8 parts of the track in the cue-poor track before (dark blue) and during (light blue) MS inactivation. (**H**) Box plots of the spatial correlation between the forth and back trials (aligned to the starting point) before and after muscimol infusion in the MS for the unidirectional cells. SH indicates the shuffle value (random pairs of cells for the back-and-forth trials). (**I**) Same than (**H**) for the non-place cells.

We then asked whether bidirectional and unidirectional place cells remaining under MS inactivation still performed distance coding. The five bidirectional place cells remaining after MS inactivation had a mean distance index of 0.92 ± 0.009, which was not significantly different from the distance index of the 38 bidirectional cells recorded before inactivation (0.89 ± 0.02, Z = 0.27, p = 0.79, two-tailed WRS test). However, while the place fields of bidirectional place cells were distributed in all parts of the track before inactivation, the place fields of place cells remaining under MS inactivation had their place fields almost exclusively near to the track ends, leaving the central part of the track not coded (Figure 4A). To quantify for this, we performed population vector overlap analysis (see Methods) (Figure 4E). While a diagonal of high correlation corresponding to distance coding was observed in all parts of the track (albeit somewhat weaker in the middle) before MS inactivation, high correlations were lost in the middle of the track and only visible at the beginning and end of the track after inactivation. Comparison with shuffled data yielded similar results (Figure 4F). We also calculated a distance overlap index (Figure 4G), which quantifies the overlap of place fields in both directions when aligned to the start of each journey. Distance overlap decreased significantly under muscimol inactivation in the center of the maze between 60 cm and 120 cm (60 to 80 cm: Z = 3.99, p < 10^4^; 80 to 100 cm: Z = 4.01, p < 10^4^; 100 to 120 cm: Z = 4.04, p < 10^4^; two-tailed WRS test). We conclude that distance coding is strongly reduced after medial septum inactivation notably in the center of the track.

These effects were not explained by the infusion procedure, as aCSF-injected animals showed no reduction in bidirectional cell number (before aCSF: 51 cells representing 33.1% of place cells; after aCSF: 53 cells representing 31.7% of place cells; Figure S6A and S6C) and no impairment in distance coding along the track (Figure S6A). Indeed, population vector analysis revealed a clear diagonal corresponding to distance coding in all parts of the track after aCSF injection (Figure S6D-E) and distance overlap analysis revealed no decrease but rather an increased overlap after aCSF injection (Figure S6F).

Given that distance coding is observed globally at the population level in the cue-poor track, including in unidirectional place cells and non-place cells, we next asked whether distance coding was also affected by MS inactivation in these cells. We performed spatial correlations between back-and-forth trials aligned to the starting point. For both unidirectional (Figure 4H) and non-place cells (Figure 4I), spatial correlations decreased significantly after the muscimol infusion (unidirectional: H(2) = 81.3, p < 10^-17^, Kruskal Wallis, before vs after p < 10^4^, multiple comparison; non-place: H(2) = 92,4, p < 10^-20^, Before vs After p < 10^8^, multiple comparison). However, for both cell populations, spatial correlations under muscimol remained significantly higher than the shuffled values (unidirectional: p = 0.0127, multiple comparison; non-place cells: p = 0.0348, multiple comparison) suggesting that some distance coding remained even after medial septum inactivation. Taken together, these results show that MS inactivation induces a global decrease in hippocampal distance coding.

To further investigate the effect of MS inactivation on hippocampal population dynamics in the cue-poor track we compared low-dimensional manifold before and after muscimol injection (Figure 5). After muscimol injection, the distance from the start of each journey less accurately mapped onto the low-dimensional manifold than before (r = 0.51 before versus 0.36 after, p = 0.165; Mann-Whitney-Wilcoxon test two-sided; Figure 5A and 5B), whereas such a tendency was not observed after aCSF injection (r = 0.38 before versus 0.39 after, p = 0.96; Mann-Whitney-Wilcoxon test two-sided; Figure 5C). However, the mapping of distance along the manifold under muscimol was significantly higher than after shuffling (see Methods) and significantly higher than the mapping of position before muscimol (Figure 5B and 5C). To decipher whether the lower mapping of distance onto the manifold under muscimol could be related to an alteration of the attractor dynamics in the network, we next compared the fraction of cell pairs showing rigid distance relationships between their firing profile across laps before and after MS inactivation. After muscimol injection, this fraction was significantly reduced (from 0.104 to 0.058, p = 0.004, two-sample Kolmogorov-Smirnov test; Figure 5D), but still significantly higher than the fraction observed with randomly shuffled spike trains (0.104 and 3.6 10-4, p < 10-8, two-sample Kolmogorov-Smirnov test; Figure 5D). Such a reduction was not observed after aCSF injections (0.104 and 0.109, p = 0.36, two-samples Kolmogorov-Smirnov test; Figure 5D).

**Figure 5:**
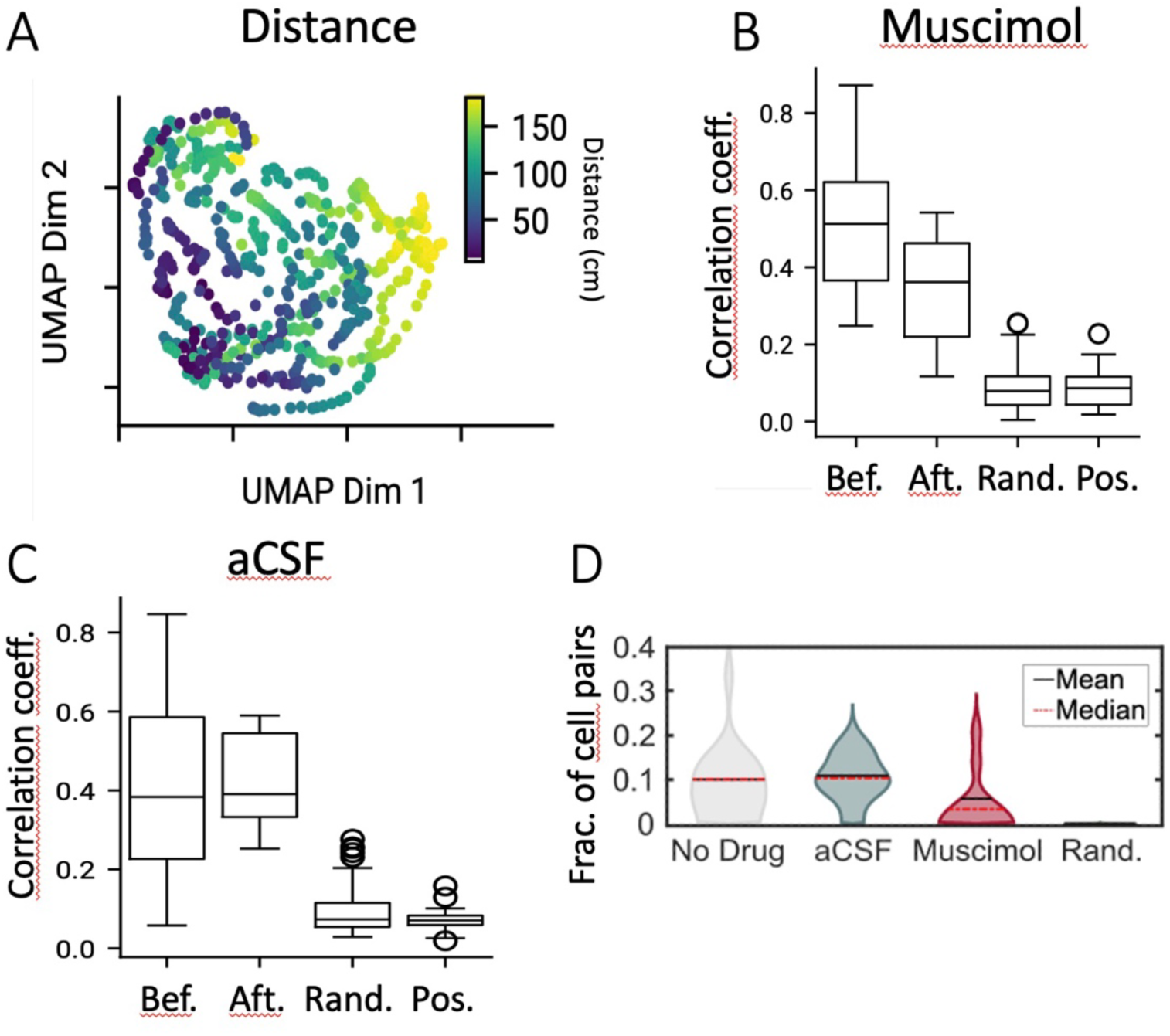
Impact of medial septum inactivation on attractor dynamics. (**A**) Example of UMAP projection after muscimol injection, colored according to distance from the start of the lap. (**B**) Correlation coefficient between UMAP projection coordinates and distance from the start of each lap before (wt) and after (inj) muscimol injection. The column (shuf) shows the correlation between UMAP coordinates and distance for lap-by-lap random shuffling of spikes. The last column shows the correlation between UMAP coordinates and absolute position in the maze. (**C**) Same as (**B**) but for aCSF injection. (**D**) Fraction of cell pairs with significant distance lag before (No Drug) and after injection of muscimol or aCSF or when cell identity is randomized.

Taken together, these results suggest that the attractor is not abolished under medial septum inactivation even if the reduced fraction of cells with rigid distance relationship suggests some alteration.

### Position coding in the cue-rich track under medial septum inactivation

Finally, we wondered whether bidirectional place cells could still perform position coding under MS inactivation. To test this, 4 mice (5 recording sessions) were exposed to a track enriched with two virtual 3D objects immediately after trials in the familiar cue-poor track under MS inactivation (Figure 6A). First, we ensured that theta power was still reduced after object inclusion. Indeed, the attenuation of theta power in this condition was 70.7 ± 14.3% (not significantly different from the 79.3 ± 4.6% attenuation before object inclusion; p = 0.81, n = 5, two-tailed WSR, data not shown). We then compared the number of bidirectional cells before and after object inclusion under MS inactivation (Figure 6A). The number of bidirectional cells decreased from 20 to 6 in the cue-poor track after muscimol injection (70% decrease), but increased to 16 after object inclusion (cue-rich under muscimol; 167% increase). Object inclusion also increased the number of bidirectional cells in a control group of 4 mice (9 recording sessions) that had received aCSF infusion in the MS (cue-rich aCSF; n = 36 before and n = 68 after, 89% increase; Figure S7A).

**Figure 6:**
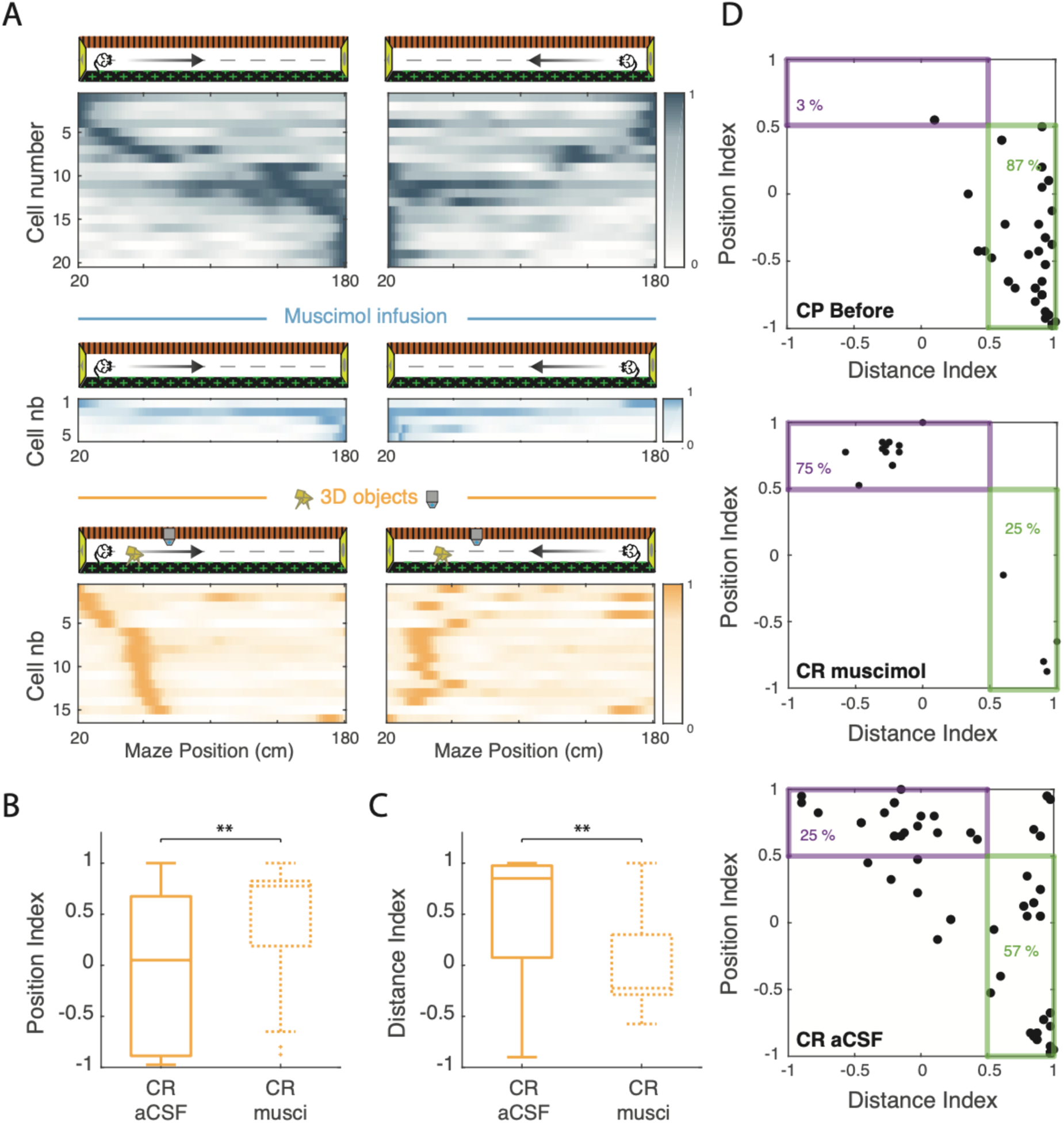
Position coding in the cue-rich track under medial septum inactivation. (**A**) Color-coded mean firing rate maps in the forth (left) and back (right) trials of bidirectional cells recorded before (top) and after the muscimol infusion in the cue-poor track (middle, blue) and in the cue-rich track (bottom, orange). The color codes for the intensity of the bin’s mean firing rate normalized on the maximal mean firing rate (peak rate) in each direction. The place cells are ordered according to the position of their peak rate in the track for all forth trials (reward zones excluded). (**B**) Box plot of the position index in bidirectional cells recorded in the cue-rich track (CR) of control mice (full orange) or muscimol mice (dotted orange). (**C**) Box plot of the distance index in bidirectional cells recorded in CR of control mice (full orange) or muscimol mice (dotted orange). (**D**) Scatter plots of distance index against position index for each bidirectional cells recorded before the muscimol infusion in the cue-poor track (CP, top), after the muscimol infusion in CP (middle) or in CR (bottom). For each track condition, cells exhibiting position or distance coding are identified by a violet or green box, respectively. The percentages of bidirectional cells showing distance or position coding are also indicated on the plots.

In the cue-rich track under muscimol most bidirectional cells had their place fields close to the two virtual objects (Figure 6A), indicating strong position coding. To quantify this, we compared the position index between the two groups (Figure 6B). The position index was significantly higher in the cue-rich track under muscimol (mean of 0.43 ± 0.16) than under aCSF (mean of −0.07 ± 0.09, Z = −2.71; p = 0.0068, two-tailed WRS test). In contrast, the distance index was significantly lower under muscimol (mean: 0.01 ± 0.13) compared to aCSF (mean: 0.54 ± 0.06, Z = 3.27; p = 0.0011, two-tailed WRS test, Figure 6C).

Finally, we analyzed the percentage of bidirectional cells coding for position or distance in the different conditions. In the cue-poor track prior to MS infusion a similar proportion of bidirectional place cells were coding for distance in both groups (aCSF: 83.3%; muscimol: 95.0%; groups (X2(2, n = 63) = 1.83, p = 0.40, chi-squared test, Figure 6D). In contrast, in the cue-rich condition under muscimol, the percentage of bidirectional cells performing position coding was much higher (75%) than in the aCSF group (24.7%) (X2(2, n = 93) = 15.6, p > 10-3, chi-squared test, Figure 6D). Accordingly, the percentage of distance coding bidirectional place cells was lower in the muscimol group (25%) than in the aCSF group (57.1%). Taken together, these results show that position coding can be observed when local cues are added to the cue-poor track under MS inactivation.

## DISCUSSION

In the course of navigation, animals can locate themselves using a set of external landmarks (position coding), but they can also use self-motion cues by integrating the distance traveled in specific directions over time (path integration-based distance coding)^1,5^. Allocentric position coding dominates the activity of place cells in real-world environments, consistent with their hypothesized role as the substrate of a cognitive map of the environment. Idiothetic coding based on self-motion cues is more likely to be observed when access to external sensory cues is reduced (when moving in darkness or away from boundaries or landmarks)^7,18,20^ or when these are unreliable^34^. Compared to allocentric position coding, idiothetic distance coding has been largely understudied, although both experimental and theoretical work suggests that it may be a prerequisite for hippocampal allocentric coding.^35^. Here, we used virtual reality for rodents, which allows better control of external sensory cues, and focused on bidirectional place cells (with place fields in both back and forth trials) to directly compare allocentric position coding and idiothetic distance coding in the same experimental paradigm.

### Sensory determinants of hippocampal distance coding

Our results show that local visual cues are necessary and sufficient for the expression of allocentric position coding in cue-rich virtual environments. Interestingly, while distance coding was expressed in both cue-rich and cue-poor environments, position coding was exclusively expressed in the cue-rich environment, specifically in the part enriched with local 3D objects. Thus, in addition to locally improving the resolution of spatial coding^26^ local visual cues can promote allocentric position coding over idiothetic distance coding. This finding complements previous studies in rats showing that distal visual cues alone are not sufficient to support allocentric position coding in virtual environments^6^. Conversely, distal cues are sufficient for allocentric position coding in real linear environments^6,27^. This difference could be due to the presence of uncontrolled local sensory cues, which are more difficult to control in real environments. Indeed, when a salient local cue (such as the wall at the end of the track) is made spatially unstable some distance coding can be observed even in real environments^34^. Alternatively, more sophisticated analytical approaches may be required to disentangle allocentric position coding from idiothetic coding in real-world settings^36^. Distance coding can also be observed when external sensory cues are decoupled from self-motion cues^37,38^ or dampened in the absence of specific task structure or reward^7^. In this later paradigm, the activity of a small fraction of hippocampal cells (∼5%) formed recurrent distance sequences spanning a fixed distance unit. This minority-based distance coding contrasts with the global distance coding we observed, in which most hippocampal neurons were involved and the sequence spanned the entire environment. The difference may be due to the fact that in our study, unlike the above study, the animals are engaged in a task with specific environmental and task-related boundaries. Further work will be needed to decipher whether this minority-based coding forms a fundamental backbone upon which the more global distance coding we observed can be built.

### Distance coding and CA1 pyramidal cells heterogeneity

CA1 pyramidal cells are morphologically and functionally heterogeneous along the dorsoventral, transverse and radial axes. Part of this heterogeneity is rooted in neurodevelopment^39,40^. This heterogeneity is best characterized along the radial axis. During spatial exploration, CA1 pyramidal cells located deep in the layer (CA1 deep, near stratum oriens) have a greater tendency to be place cells within and across environments, whereas CA1 sup cells tend to fire more sparsely in a context-specific manner^41,42^. Furthermore, CA1deep cells tend to be active in cue-rich parts of heterogeneous environments while CA1sup cells tend to be active in cue-poor parts away from landmarks^11,20^. Consistent with this, we observed an increased propensity of CA1deep cells to perform position coding and CA1sup cells to perform distance coding in our heterogeneous cue-rich environment. In the cue-poor track, however, both CA1deep and CA1sup cells performed distance coding in similar proportions, with similarly high distance coding indices. Furthermore, the proportion of place cells among active cells did not differ between CA1deep and CA1sup cells in the cue-poor environment.

These results show for the first time that, despite important differences in morpho-functional properties and connectivity, both CA1deep and CA1sup cells can be equally recruited to code distance in cue-poor environments. These results seem a priori inconsistent with previous work showing that CA1deep cells behave as landmark vector cells, systematically discharging next to external landmarks on a cued treadmill, whereas CA1sup cells can be activated away from external landmarks^32^. According to this work, CA1deep cells would be primarily driven by external sensory cues, whereas CA1sup cells would be preferentially driven by self-motion or idiothetic cues. However, in our study, both cell types were equally recruited in the cue-poor track. One possibility is that both cell types perform distance coding, but on the basis of different sensory cues. While CA1deep cells would code distance with respect to an external landmark, such as the end of the track, CA1sup cells would use self-motion or idiothetic cues to code distance based on path integration. However, if this was the case, CA1deep cells would preferentially code at the ends of the path where a cue is visible, whereas CA1sup cells would preferentially code at the beginning of the path before error accumulation would prevent distance coding based on path integration^43^. However, CA1deep cells were active from the beginning of the track, while CA1sup cells were also active at the end of the track. Alternatively, both CA1deep and CA1sup cells could be activated by default based on self-motion cues, but when present, external sensory cues could take control over CA1deep cells (but not CA1sup cells), favoring allocentric position coding. How distance and position coding map onto cellular heterogeneity in other hippocampal areas remains to be investigated^44^.

### Attractor dynamics of hippocampal distance coding

Contrary to place field analysis, dimensionality reduction does not explicitly refer to behavioral variable like absolute distance or position. In particular, it allows highlighting relative distance coding along the track. Dimensionality reduction analyses applied to hippocampal coding at the population level in the cue-poor track revealed a 1D manifold. Distance traveled along the track, but not position, mapped well onto this manifold. Indeed, there was a good correlation between the evolution of population activity along the manifold and the distance traveled by the animal along the maze. The observation of this manifold suggests that a Continuous Attractor Network (CAN) might constrain hippocampal activity. The population activity of large ensembles of grid cells has recently been shown to also lie on a toroidal (2D) manifold^14^ imposing rigidity in the correlation structure of their activity^12,33^. We found a similarly fixed distance relationship between hippocampal cell activity in this condition, suggesting that grid cell attractor dynamics may constrain hippocampal activity in the absence of local visual cues. Indeed, grid cells discharge at fixed distances in 1D environments^15,45^ and are more influenced by self-motion-related information than visual information during gain changes in virtual environments^46,47^. Consistent with this hypothesis, MS inactivation, which disrupts the spatial organization of grid cell discharge^16,17^ and distance estimation based on path integration ^19^ strongly reduced distance coding in the absence of local visual cues, with a significant reduction in the number of bidirectional distance-coding cells. Population vector analysis further confirmed a significant loss of distance overlap, particularly in the center of the path. These deficits were not observed when aCSF was injected into the medial septum as a control. Accordingly, we observed a decrease in the correlation between the evolution of population activity along the manifold and the distance traveled by the animal along the maze, together with a loss of rigidity in the distance relationship between place fields across laps. It is worth noting that although distance mapping along the manifold and correlation structure of hippocampal population activity were reduced under MS inactivation, they were still higher than when spiking was shuffled. This suggests that the underlying CAN structure may be preserved. Interestingly, similar differences between coding in physical and manifold space have been observed for grid cells^14^. For example, grid cells retain some distance coding properties and the correlation structure of their activity is preserved even when the grid pattern is lost in the dark^23,24^ or after hippocampal inactivation^48^. Collectively, these results suggest that grid cell activity could sustain hippocampal distance coding in the absence of local visual cues and that MS inactivation would more specifically alter the mapping of distance onto the manifold rather than the underlying attractor dynamics. The few bidirectional distance-coding cells that remained under MS inactivation were active at the start or end of the track, suggesting that they used allothetic cues (the track ends) rather than idiothetic cues to encode distance. This intuition was confirmed by population vector analysis, which showed that some distance overlap remained at the start and end of the track. These end-of-track cues could be used to anchor the grid in the control condition, allowing accurate path integration-based navigation and hippocampal distance coding^49,50^. Such anchoring could be lost under MS inactivation thus precluding accurate distance coding along the track while potentially preserving path-integrated distance coding along the manifold.

### MS inactivation alters distance but not position coding

Allocentric position coding could be observed under MS inactivation when local visual cues (virtual 3D objects) were introduced into the cue-poor track. In this condition, position coding cells were concentrated in the cue-rich part of the track and dominated hippocampal spatial coding. This is consistent with behavioral results showing that grid cell anchoring to the track is not required for cue-based navigation^19,49^. Previous studies have reported various effects of medial septum inactivation on place cell coding in real-world environments. While allocentric position coding was observed in small scale 2D and 1D environments under MS inactivation^17,18^, hippocampal coding was selectively altered when animals ran in a wheel during a spatial working memory task or away from external cues sensory cues^18,20,51^. However, allocentric and idiothetic coding have been difficult to disentangle in these experiments. Our experimental paradigm using back-and-forth trials in virtual reality together with local cue manipulations allowed for the first time to disentangle the effect of MS inactivation on allocentric position coding and idiothetic distance coding. Our results are consistent with a specific deficit in hippocampal distance coding based on path integration but not allocentric position coding. As the MS comprises a variety of cell types, future work should determine which of these cells are more specifically involved in hippocampal distance coding. In addition to altering grid cell firing, the effects of MS inactivation on hippocampal distance coding could result from altered theta rhythmicity and theta-driven dynamics of internally organized hippocampal networks, such as theta sequences. Indeed, a computer model has shown that self-motion-based firing fields can be built from theta sequences in the absence of external sensory cues^18^. However, we and others have previously shown that theta phase precession and theta phase spike coordination are altered in the cue-poor condition even without MS septal inactivation^11,26^. Collectively, these results suggest that theta sequences are dispensable for distance-coding firing fields, but may be selectively involved in internally generated firing fields in the context of working memory. Finally, while average trial speed was not significantly affected by MS inactivation, the number of stops per trial was increased after compared to before MS inactivation. This could be due to direct inhibition of MS glutamatergic neurons, whose activity has been directly linked to locomotor speed^52^. Part of this increase could also be related to a decrease in motivation for reward at late stages of the recording session when spatial coding is assessed under MS inactivation. In line with this, the number of stops was also increased at similarly late stages of the experimental protocol after control aCSF injections. Although the impact of behavior on hippocampal dynamics is difficult to disentangle from the impact of hippocampal dynamics on behavior, the increased number of stops following aCSF injections did not prevent clear distance coding in this condition. Conversely, altered hippocampal distance coding and path integration-based self-localization deficits could also contribute to the increased number of stops observed after MS inactivation.

### Conclusions

Despite years of investigation, the specific determinants of hippocampal distance coding are unclear^1^. Using virtual reality and local cue manipulations, we could isolate distance and investigate its cellular and network determinants. We showed that distance coding is ubiquitous in the hippocampus in the absence of local visual cues, involving both CA1deep and CA1sup cells as well as unidirectional and non-place cells. In this condition, the distance traveled along the track, but not the allocentric position, could be mapped onto a low-dimensional manifold imposing rigid distance relationships between place fields. This is reminiscent of attractor dynamics observed in the grid cell network. MS inactivation, which alter the firing pattern of grid cells, dampened distance coding at the single cell level and rigid distance relationships at the population level. However, allocentric position coding could still be observed under MS inactivation in the presence of local visual cues. Based on these results, we propose a specific role for grid cells in maintaining hippocampal distance coding in cue impoverished conditions.

## METHODS

### Animals

Data were collected from 16 male mice C57BL/6J (Janvier/Charles River) between 8 and 12 weeks of age during the recording phase (weight: 21-23.6 g). Mice were housed in groups of two or three per cage prior to the first surgery and then individually housed on a 12-hour inverted light/dark cycle. Training and recording were performed during the dark phase.

### Ethics

All experiments were approved by the Animal Care and Use Committee of the Institut National de la Santé et de la Recherche Médicale (INSERM) and authorized by the Ministère de l’Education Nationale de l’Enseignement Supérieur et de la Recherche after evaluation by a local ethics committee (agreement number 02048.02), in accordance with the European Community Council Directives (2010/63/UE).

### Surgical procedure to prepare head fixation and cannula implantation

Animals were maintained under isoflurane anesthesia supplemented with a subcutaneous injection of buprenorphine (0.1 mg/kg) throughout the surgical procedure. Two jeweler’s screws were inserted into the skull above the cerebellum for reference and grounding. For muscimol infusion in the MS, a guide cannula (26G, Dominique Dutscher) was implanted 1 mm above the MS (AP: +1 mm, ML: +0.7 mm, DV: −3 mm, angle 10° towards the midline). A dental cement cap was then constructed, leaving the skull free above the hippocampi for later craniotomies. The exposed skull was sealed with silicone elastomer (Kwik-Cast, World Precision Instruments). A small titanium rod (0.65 g; 12 x 6 mm) was inserted above the cerebellum to serve as a fixation point for a larger head plate, which was used for head fixation only during training and recording. After surgery, buprenorphine (0.1 mg/kg) was administered twice daily for 3 days. Mice were allowed to recover for 5 days before behavioral training began.

### Virtual reality set up

A commercially available virtual reality system (Phenosys Jetball TFT) was combined with a custom-designed, 3D-printed, concave plastic wheel (center diameter: 12.5 cm; side diameter: 7.5 cm; width: 14 cm, covered with a white silicon-based coating) to provide 1D motion with a 1/1 coupling between the movement of the mouse on the wheel and the movement of its avatar in the virtual reality environment. The wheel was surrounded by six 19” TFT monitors covering a total angle of 270 degrees. The monitors were elevated so that the mouse’s eye level corresponded to the bottom third of the screen height to account for the fact that the rodent’s field of view is upwardly biased. The head restraint system (Luigs and Neumann) was placed behind the animal so as not to interfere with the view of the virtual reality environment. The virtual reality environment was a 200 cm long and 32 cm wide virtual linear maze with different patterns on the side and end walls, enriched or not with virtual 3D objects (see Virtual Reality Environments section). The movement of the wheel updated the position of the mouse avatar. The mouse could only move forward or backward, but could not turn back in the middle of the path (see Training section).

### Virtual reality environments

#### Cue-poor track

Each side wall had a unique pattern (black and orange stripes on one wall; green crosses on a black background on the other). End walls had gray triangular or round shapes on a yellow background.

#### Cue-rich track

This maze was identical to the cue-poor track in terms of wall patterns and dimensions, but two or three virtual objects were placed on the sides between the animal’s path and the walls. The objects were a yellow origami crane (dimensions: 9 x 9 x 7 cm; position: 37 cm from end wall), a blue and gray cube (dimensions: 5 x 5 x 5 cm; position: 64 cm from end wall), and a tree (15 x 15 x 22 cm; position: 175 cm from end wall). The animal could neither orient to the objects nor receive any sensory feedback from them other than vision. 3 mice used in this study (6 recording sessions) were trained and recorded in a virtual linear track enriched with 2 virtual objects (origami + cube). An additional mouse trained and recorded in a track enriched with 3 objects was added to the study. The results of this animal were already used in a previous study (Bourboulou, Marti et al., 2019). For the experiments in which animals were first trained in the track without objects and then exposed to a novel environment with objects, the track with 2 objects was used.

### Training

Mice were first habituated to the experimenter by daily handling sessions of 20 min or more, which continued throughout the experiment. After a 5-day recovery period following surgery, mice were water-deprived (1 ml/day, including the amount of water consumed during training). They were then progressively trained to run in the virtual reality setup. First, mice were familiarized with running head-fixed on the wheel for water rewards in a black track (screens always black). During these sessions, animals received sweetened water (5% sucrose) as a reward for every 50 cm run on the wheel. When the animals were familiar with the setup, they were trained to run in a 200 cm long linear virtual track (familiar track). When animals reached the end of the track, a liquid reward delivery tube extended in front of the animal and the animal had to lick to obtain the reward (a 4 mL drop of water containing 5% sucrose). Animals were then teleported to the same position but in the opposite direction of the maze and had to run in the opposite direction to the end of the maze to receive another reward. Animals were trained until they were able to complete at least 60 trials. Ad libitum access to water was restored if the animal’s weight dropped below 80% of its preoperative weight at any time during training.

### Recording procedure

When animals reached a stable behavioral performance (at least two reward/minute and 60 trials), we performed acute recordings using silicon probes (4/8 shanks; A-32/A-64 Buzsaki Probe, Neuronexus). The day before the first recording session, animals were anesthetized (with isoflurane supplemented with buprenorphine 0.1 mg/kg) and a craniotomy was made over one hippocampus (centered at a point AP: −2 mm ML: ± 2 mm). The craniotomy was covered with agarose (2% in physiological Ringer solution) then sealed with silicon elastomer (Kwik-Cast, World Precision Instruments). This craniotomy was used for acute recording for 2-3 consecutive days (with the probe lowered to a new location each time). A second craniotomy was then made over the other hippocampus using the same procedure, and recordings were made for 2-3 additional days. The silicon probe was lowered into the brain while the animal was allowed to walk freely on the wheel with the screens displaying a black background. The good positioning of the probe with recording sites in the CA1 pyramidal cell layer was verified by the presence of multiple units showing complex spike bursts at several recording sites and the recording of sharp wave ripples during quiet behavior. After positioning the silicon probe, the virtual reality environment was displayed on the screen. On the day of the last recording in each hippocampus, the back of the probe shaft was coated with a thin layer of a cell-labeling red fluorescent dye (DiI, Life technologies), so that its location (shaft tips) could be assessed histologically post hoc. All mice (n = 16) were recorded in a familiar environment (either cue-poor or cue-rich track) for approximately 20 trials. For mice trained in the cue-poor track (n = 10), these trials were followed by an intracranial infusion of muscimol (n = 5) or aCSF (n = 6) in the MS (see ‘Intracranial infusions’ section), during which the mouse was free to run with the screens turned off (black). Approximately 15 min after the infusion, mice were again exposed to the familiar track for at least 20 trials. Some mice (n = 4 for muscimol, n = 4 for aCSF) were then exposed, 3 min after the end of the second session, to a new environment (cue-rich track), identical to the previous one except for the presence of two 3D objects (an origami and a cube). Again, animals had to complete at least 20 trials in this last condition. Note that animals stayed head-fixed on the wheel surrounded by screens during the entire recording session.

### Intracranial infusions

Intracranial infusions were performed as follows. First, a custom-made infusion cannula (33 Ga, Dominique Dutscher) was connected by polyethylene tubing to a 2 µl Hamilton syringe mounted in a microinjection pump (Legato 130 Single Syringe I/W Nanoliter, World Precision Instruments). The syringe, tubing, and cannula were filled with pure water. Dissolved muscimol or aCSF was then added to the infusion cannula (separated from the water by an air bubble). Muscimol powder (Sigma-Aldrich) was dissolved in sterile aCSF (Harvard Bioscience, Inc.) at a concentration of 1 µg/µl. At the beginning of the microinjection procedure, the infusion cannula was inserted into the implanted guide cannula and the infusion was made 1 mm deeper than the tip of the guide cannula. 0.1 µl of muscimol or aCSF was slowly injected into the MS (0.08 µl/min), and the infusion cannula was left in place for another 2 min after microinjection was completed. During infusion, the animals were allowed to run freely and were given food. Animals showed no signs of stress or discomfort during the procedure. 10-15 min after microinjection, animals were returned to the virtual familiar environment.

### Data acquisition and pre-processing

The animal’s position in the virtual maze was digitized by the virtual reality control computer (Phenosys) and then sent to a digital-to-analog card (0-4.5V, National Instrument Board NI USB-6008) connected to the external board (I/O Board, Open Ephys) of a 256 channel acquisition board (Open Ephys). Neurophysiological signals were acquired continuously at 25 000 Hz on a 256-channel recording system (Open Ephys, Intan Technologies, RHD2132 amplifier board with RHD2000 USB interface board). Spike sorting was performed semi-automatically using KlustaKwik (Rossant et al., 2016; https://github.com/klusta-team/klustakwik). Clusters were then manually refined using cluster quality assessment, auto- and cross-correlograms, cluster waveforms, and similarity matrix (Klustaviewa, Rossant et al., 2016).

### Data analysis

Data analysis was performed using custom written programs in using MATLAB or python softwares.

### Reward zones definition

The reward zones, located between the maze extremities and 10% of the track length (0–20 cm and 180–200 cm), were not considered in the analysis.

### LFP Theta attenuation analysis

The original broadband signals were low-pass filtered (0-500 Hz) and downsampled to 1250 Hz for local field potentials. The time-frequency spectrogram (0-20 Hz) of the LFP was computed using a window size of 5 s and a time step of 1 s, and was 1/f corrected. The LFP was band-pass filtered in the delta (0.5 - 4 Hz) and theta (6 - 9 Hz) bands using zero-phase IIR Butterworth filters. Theta power was computed when the animal was performing the task in each track condition before and after the infusion. We first determined the theta power by Hilbert transform and calculated the theta power on the channel with the best ratio of theta to delta power when the animal speed exceeded 2 cm/s. Theta power was taken from the maximum power in the 6-9 Hz band. To examine changes in theta power after infusion of aCSF or muscimol in different recording sessions, the absolute theta power was normalized by dividing by the total power of the band (1-30 Hz). This normalization allowed comparison of pre- and postinfusion theta power levels across recording sessions. A second method was used to calculate the theta power in each movement period. Movement periods were detected using criteria for running speed and duration. Specifically, a locomotor period was defined as a period of at least 2 seconds with a velocity greater than 2 cm/s. Two movement periods were merged if less than 0.5 sec separated them. The theta power in the movement period was averaged. For both theta power methods, the percentage of theta attenuation was computed from the ratio of theta power after the infusion to the theta power before the infusion.

### Speed and stop analysis

The maze was divided into 100 spatial bins of 2 cm, and the mean speed was calculated for each bin. The mean speed before or after muscimol infusion was the average of all bin speeds in each condition. We considered that the animal was making a stop if its speed was less than 1 cm/sec in a spatial bin (at least 2 seconds stop).

### Firing rate map

The maze was divided into 100 spatial bins of 2 cm. For each trial, the number of spikes and the occupancy time of the animal in each spatial bin were calculated to obtain the number of spikes vector and the occupancy time vector, respectively. These vectors were smoothed using a Gaussian filter with a half-width set to 10 spatial bins. Spikes occurring during epochs when the velocity was less than 2 cm/s were removed from all analyses. The smoothed spike count vector was divided by the smoothed occupancy time vector to obtain the firing rate vector for each trial. Firing rate vectors were pooled for a given condition (track condition, pre- or post-infusion) and animal direction (e.g., back) to create a firing rate map. These pooled vectors were also averaged to produce the mean firing rate vector, which corresponds to the mean firing rate for each spatial bin.

### Pyramidal cell classification

Cells with a mean firing rate lower than 20 Hz and either a burst index greater than 0 or the spike duration greater than 0.4 ms were classified as putative pyramidal neurons ^53^. They were classified as interneurons otherwise. To distinguish deep pyramidal cells from superficial pyramidal cells, we defined the middle of the CA1 pyramidal layer as the recording site where ripples had the largest power (Geiller et al., 2017; Mizuseki et al., 2011; Sharif et al., 2021). To detect ripple events, a spectrogram (using a short-time Fourier transform) of the LFPs in CA1 pyramidal layer was computed (time windows of 20 ms) for one channel (the one with the strongest ripple power between 120 and 230 Hz) of each shank. Ripple epochs were defined as periods during which ripple power (120 to 230Hz) was continuously greater than mean 120 to 230Hz power + 3 standard deviation (sd) and than mean 250-300Hz + 3 sd. Then, during ripples times, the power in the ripple band was calculated for each channel and the channel with the largest mean power was selected as the middle of the pyramidal layer (Mizuseki et al., 2011). To locate the approximate location of the cell body, we used the method described in Geiller et al., 2017.

### Active cells classification

A cell was considered active if the mean firing rate was greater than 0.3 Hz, the peak firing rate was greater than 1 Hz, and the cell fired at least one spike in 50% of the trials. These three criteria had to be verified in either the forth or back direction. The percentage of active cells in each recording session was calculated from the total number of pyramidal cells. All recording sessions had at least 15 recorded neurons.

### Place cells classification

A place cell was defined as a pyramidal cell that showed a mean place field in at least one direction. A bootstrap procedure was used to detect a mean place field. For each trial, a new spike train was generated using a Poisson process with λ equal to the mean firing rate of the trial and a time interval of 1 ms. A “randomized” firing rate map was then generated and the mean firing rate vector was determined and compared to the mean firing rate vector from the initial rate map. This process was repeated 1000 times to generate a p-value vector (p-value for each 2 cm spatial bin). Candidate place fields were defined as a set of more than three contiguous spatial bins associated with p-values less than 0.01Two place fields were merged if the distance between their closest edges was at most equal to five spatial bins (10 cm). The edges of place fields were extended by a maximum of five spatial bins (for each edge) if the p-value for these bins was less than 0.30. A field with a size greater than 45 spatial bins (90 cm) was not considered as a place field. To validate a mean place field, the cell had to verify a stability criterion. Spatial correlations were calculated between the firing rate vector of each trial and the mean firing rate vector. The spatial bins corresponding to other detected place fields were not included in the spatial correlations. The place field was validated if the spatial correlations were greater than 0.60 in at least 40% of the trials. If multiple mean place fields were detected, only the place field with the highest peak was retained, unless otherwise specified. An active cell with at least one place field in one direction was considered a unidirectional place cell. Bidirectional cells were place cells with at least one place field in both directions. The proportion of place cells was calculated from the total number of active cells. All recording sessions had at least 13 active cells. To calculate the proportion of bidirectional and unidirectional cells, only sessions with at least 4 active cells were included. The same procedure was used to calculate place fields per lap without the stability criterion, which cannot be calculated on single trials. A place field per lap was conserved if it overlapped at least one spatial bin with the closest mean place field.

### % of bidirectional cells and relative change

To quantify the effect of muscimol infusion on the number of bidirectional place cells, we calculated the percentage of bidirectional place cells on place cells before and after muscimol infusion. Only sessions with at least 4 place cells were kept for this analysis. To quantify the effect of MS inactivation on each cell category (unidirectional, bidirectional, or non-place cells), the relative change was calculated by dividing the difference in the number of cells (after-before) by the number of cells before the MS infusion.

### Position and distance index

For each bidirectional cell, a position and distance index was calculated using the mean firing rate maps in both directions. To calculate the distance index, the mean firing rate maps in both directions were aligned to the starting point. The distance from the starting point to the peak rate of the place field was calculated for each direction. This formula was then used to calculate the distance index (DI):

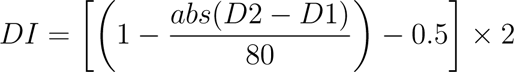

where abs(D2-D1) is the absolute value of the distance in bins between the peak rate positions in both directions. 80 is the number of bins (excluding reward zones). If there was complete overlap between the place fields, the distance index would be one. If there was a difference of 40 bins between them, the index would be equal to 0. Position indices were computed using the same formula, except that the firing rate maps were aligned to the external reference frame. Position index with values close to −1 correspond to distance coding cells with place fields at the ends of the track. Bidirectional cells with a position index and a distance index both greater than 0.5 were excluded from the analysis, as these cells had a place field in the middle of the trace and it was not possible to disentangle position and distance coding in this case (3 out of 90 in the cue-poor track and 6 out of 99 in the cue-rich track).

### Position and distance-coding cells

Cells with a position index greater than 0.5 and a distance index less than 0.5 were classified as position-coding cells. Conversely, cells with a distance index greater than 0.5 and a position index less than 0.5 were classified as distance-coding cells. In these two categories of cells, the distribution of place fields between the cue-rich and cue-poor parts was examined by comparing the proportion of place fields located in a 40 cm zone enriched with local visual objects with the proportion of place fields located in a 40 cm zone deprived of cues.

### Population vector analysis

To assess the quality of distance coding at each position in the track, a population vector analysis was performed using the place field maps of both directions. The place field maps indicate the presence (1) or absence (0) of a place field for all bidirectional cells. Accordingly, each bin of the place field map contained a vector indicating the presence or absence of a place field for each bidirectional cell. We then correlated (Pearson’s correlations) the vectors from the place field map of the forward trials with all the vectors from the place field map of the backward trials to obtain an 80*80 correlation matrix. For statistical comparison, we first generated 500 randomized firing rate maps for each cell (using the same method as for place field detection) and computed the population vector analysis for each randomized firing rate map. We then looked for correlations in the real data correlation matrix that were above the 99% percentile calculated from the randomized correlation matrices. These correlations were considered significant.

### Distance overlap

Another method used to assess the quality of distance coding at each location in the track was distance overlap. First, place field maps of both directions were aligned to the starting point, and for each bin we computed the ratio of the number of cells coding for distance (showing place field overlap) to the total number of bidirectional cells coding for distance in at least one direction for that position. This value measures the probability, at each position bin, that bidirectional place cells are coding for the distance. For better comparison between conditions (pre- vs. post-infusion), we averaged the distance overlap values in bins of 20 cm.

### Spatial correlation

To assess distance coding in unidirectional cells (cells that have a place field in only one direction) and non-place cells (cells that have no place field) we performed, for each cell, spatial correlation between the mean firing rate vector in one direction and the mean firing rate vector in the other direction, with both firing rate vectors aligned to the starting point. These values were compared with shuffling values corresponding to spatial correlation between random pairs of cells in the back and forth trials.

### Stability index

The stability index of a cell was computed as the mean of the spatial correlations between all pairs of firing rate vectors. This way, the cell stability index takes into account the activity patterns of all trials and provides a reliable quantification of the inter-trial reproducibility of the cell’s activity. Note that this stability index differs from common stability indices based on correlations of mean firing rates between even and odd trials or between two halves of the same recording session, so the values obtained are not directly comparable.

### Spatial Information

The spatial information (SI) was calculated according to the following formula (Skaggs et al., 1996):

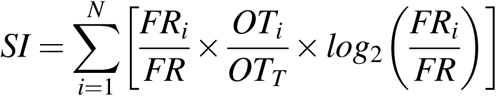

where N is the number of spatial bins (N = 100), FR_i_ is the mean firing rate determined in the i-th spatial bin, FR is the mean firing rate, OT_i_ is the mean occupancy time determined in the i-th spatial bin, OT_T_ is the total occupancy time based on the mean occupancy time vector.

### Out/in-field firing ratio

The out/in-field firing ratio was computed as the ratio of the mean firing rate outside the mean place field (excluding secondary place fields) to the mean firing rate inside the mean place field.

### Place field dispersion

A place field dispersion measure has been computed to quantify how much each place field per lap was dispersed around the mean place field. The place field dispersion (PFD) was calculated using the following formula:

where C is the center of the mean place field, Ci is the center of the field in the i-th lap and M is the number of laps with a single-trial detected field, L is the total length of the maze and N is the number of spatial bins. The center of a place field was defined as the spatial bin with the highest firing rate.

### Place field width

Place field width was computed as the distance between the place field edges and only determined for entire place fields. A place field was considered as complete when its firing rate increased above 30% of the difference between highest and lowest place field activity and then dropped below this threshold.

### Statistics

All statistical analyses were performed using MATLAB codes (MathWorks). For each distribution, a Lilliefors goodness-of-fit test was used to test whether the data were normally distributed, and a Levene test was used to test for equal variance. To compare two independent distributions, the unpaired Student t test was used if normality or equal variance was verified, otherwise the Wilcoxon rank sum test was used. Wilcoxon sign rank or paired t-test was used to compare paired samples. For multiple comparisons, 2 independent factors ANOVA was used to test for normality and equal variance between samples, otherwise Kruskal-Wallis test with Bonferroni post hoc test was used. The chi-squared test was used to compare the percentages of cell category (unidirectional vs bidirectional or for bidirectional cells only: distance coding VS position coding VS unclassified) between track condition (cue-rich and cue-poor) or between infusion condition (before vs after or muscimol vs aCSF).

### Spatial cross correlation analysis

For the cross-correlation analysis, the time-course of activity for each cell was split into laps within the same condition. The 200 cm track was divided into *n* = 100 bins of equal size. Firing rates were calculated as the number of spikes fired in a spatial bin divided by the time spent by the animal in it. Only the data points in which the rodent had a velocity higher than 2*cm/s* were included in the binning procedure. Firing rates were smoothed with a sliding Gaussian window and with a half width of 10 bins.

To extract the spatial shift between pairs of cells we first calculated, with a custom MATLAB function, the two-point correlation 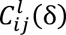 between the firing rates 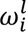 and 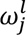 for the cell pair (*i*, *j*) in lap *l* and at a spatial lag δ, namely:

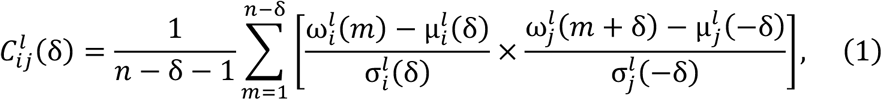

where *n* is the number of spatial bins, δ is the lag defined as a discrete number of spatial bins, µ and σ are, respectively, the mean and the standard deviation of the firing rate 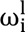 defined as follows:

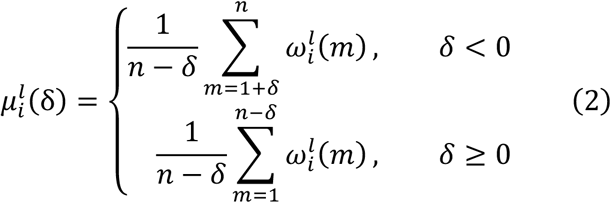

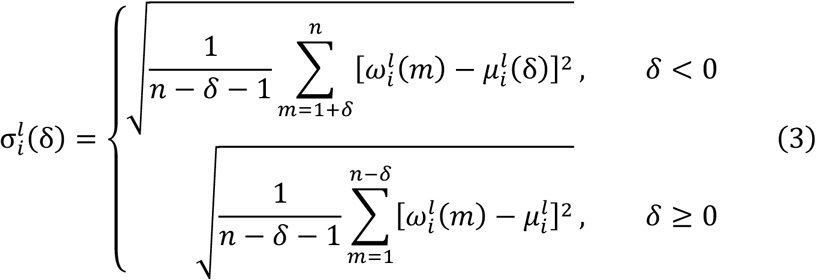

For each pair of cells, we considered both positive and negative lags by means of the following equivalence:

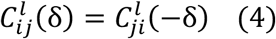

The maximum lag considered was *δ_max_* = 70, corresponding to a physical distance of 140cm. The order of spatial bins in backward laps was reversed to measure the distance from the start of the lap rather than the absolute position on the track.

Finally, the cross-correlation function for the (*i*, *j*) cell pair was obtained by averaging out the lap dependence:

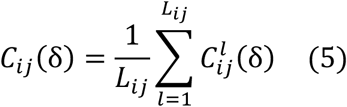

where *L_ij_* is the number of laps in which both cells of the pair were active. To avoid small sample errors, we included only the cell pairs for which *L_ij_* was higher than 80% of the total number of laps in the condition considered. We call *N*_80%_ the number of such pairs.

To assess the significance of each cross-correlation value *C_ij_*(*δ*) we compared it with a distribution Γ*_ij_*(*δ*) obtained from a set of 1000 randomizations of the dataset. Each randomization was performed on the spike counts arrays for the (*i*, *j*) cells by reshuffling the order of the spatial bins for each cell and lap. The cross-correlation procedure was then performed on the randomized dataset, after smoothing, to obtain a correlation value for each cell pair and lag *δ*. A *p-value p_ij_*(*δ*) was assigned to each cross-correlation value *C_ij_*(*δ*) through the *z-score* derived from the comparison with the Γ*_ij_*(*δ*) distribution. Among all possible pairs (*i*, *j*), we selected as significantly correlated only those with a cluster *S* of at least *S_cluster_* = 10 adjacent lags *δ* for which *p_ij_*(*δ*) < 0.001. The fraction of such pairs was then calculated by dividing the number of selected pairs by *N*_80%_.

The spatial shift *D_ij_* in the activations of the cell pair (*i*, *j*) was then chosen to be the distance value belonging to a significant cluster and corresponding to the peak of cross-correlation, namely 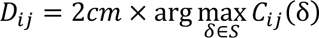. For each mouse and condition, a distance matrix was created by the collecting the values |*D_ij_*| from all selected pairs. The multidimensional scaling on such matrices was then performed with the MATLAB *mdscale* function using the *metricstress* criterion and a number of replicates equal to 50.

### UMAP Analysis

#### 1. Population Activity Analysis

##### 1.1. Preprocessing

- Spike Train Convolution: The spike trains were convolved with a 200ms Gaussian kernel to smooth the binary spike trains into a continuous representation of the neural firing rate over time.
- Subsampling: The convolved signal was subsampled down to 5Hz to reduce the dimensionality of the data while preserving the low-frequency components relevant for population activity analysis.
- Velocity Thresholding: Time points where the animal’s velocity fell below 2cm/s were excluded from the analysis to remove periods of inactivity where neural activity might not lie on the 1D manifold.
- High Firing Rate Neuron Exclusion: Neuronal clusters with a basal firing rate exceeding 8Hz, determined by a threshold based on the population firing rate histogram, were excluded to minimize the influence of potential interneurons on the UMAP dimensionality reduction.
- Time window selection: session time limits where sometimes adjusted to account for probe movements hightlighted by raster plot analysis.

##### 1.2. Dimensionality Reduction with UMAP

UMAP (Uniform Manifold Approximation and Projection) [1] was used to reduce the dimension-ality of the preprocessed population activity data. UMAP preserves the underlying distance relationships between data points in a lower-dimensional space, making it ideal for exploring the structure of neural activity patterns.

UMAP was configured with the following parameters:

- Target Dimensionality: 2 (embedding the data into a two-dimensional space)
- Number of Neighbors: 30 (specifying the number of neighboring data points used in the local neighborhood structure)

The resulting two-dimensional embedding allows for visualization and further analysis of the population activity patterns across the recorded neural clusters.

##### 1.3. Analysis of Tuning Curves

- Identifying 1D Manifold Modulated Neurons: To identify neurons whose activity is modulated by the 1D manifold state, tuning curves were extracted from the UMAP-embedded data.
- Curvilinear Abscissa: A spline function was fit to the two-dimensional embedding to obtain a coordinate along the 1D manifold. This curvilinear abscissa reflects the continuous nature of the coding manifold and the animal’s exploration path. The procedure consists first in an automatic fit, and second by a human-controlled verification and correction. The steps for the automatic procedure are the following: i. K-means clustering wih 4 clusters was performed on the UMAP positions of all data points. ii. Hierarchical clustering was used to order correctly the k-means centroids along the presumed trajectory. iii. A B-spline curve was interpolated through these centroids to define the curvilinear abscissa.
- Human-controlled verification and correction. The automatically generated curvilinear abscissa was then manually refined if necessary to better capture the overall structure of the UMAP data by refining the position of the centroids.
- Tuning curve Construction: Tuning curves were built similarly to place cell tuning curve but using the curvilinear abscissa as parameter instead of the raw spatial coordinate of the animal. Each tuning curve consisted in 50 bins spanning the entire extent of the curvilinear abscissa.
- Statistical significance: a standard statistical test for place field significance [2–5] was employed to assess the statistical significance of theses tuning curves. A circular shift validation method was used. For each lap independently, the spike trains were shifted in time circularly after removing immobility periods. Tuning curves were then generated for the shifted spike trains. The original tuning curve was considered significant if its maximum value exceeded the 5th percentile of the maximum values obtained from the shifted controls. Only tuning curves surpassing this significance threshold were retained for further analysis.

##### 1.4. Analysis of the correlation between the UMAP projection and spatial coordinates

To assess whether the population of neurons encodes for absolute position or distance, a correlation coefficient between the UMAP projection and the corresponding spatial coordinates for each dataset was computed.

To achieve this, we denote the coordinates of the *i*-th projected population activity vector as (*xi*, *yi*) and the corresponding spatial coordinate *si*. We then center the data 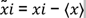 and 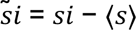, where ⟨.⟩ denote the mean value over the whole set of points. We then define the correlation coefficient as:

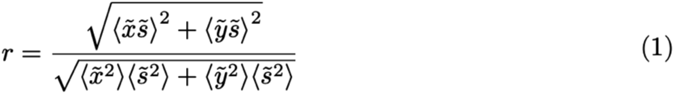

This coefficient ranges from 0 to 1. It essentially represents the normalized norm of the electric dipole moment for the distribution of charges *si* at position (*xi*, *yi*). A value of *r* = 1 indicates a perfect linear relationship between the UMAP projection and the spatial variable *s*. Conversely, a value close to 0 suggests a symmetry *s* ↔ −*s* in the UMAP projection, or a global mixing. Moderate value can be achieved in the case of a cloud such that one of its axis correlates with *s*. The larger the extension along the orthogonal axis, the smaller the value of *r* will be.

##### 1.5. Global statistics

To assess the dominant coding scheme, we calculate the set of correlation coefficients *r* between the UMAP projection and the distance with respect to the start of the lap, or between the UMAP projection and the absolute position. To assess the significance of our values, we compare it with those obtained after temporal circular shift of the spike trains (see above). The comparison between both set of *r*-coefficients highlights the distance coding scheme, the correlation with the position being comparable with the value after circular shift.

To examine the impact of muscimol infusion, we calculate the set of *r* correlation coefficients between the UMAP projection and the distance with respect to the start of the lap (i) in the WT condition, (ii) after injection (of muscimol or aCSF).

##### 1.6. Source code

For the population activity analysis, scripts are written in Python and available at https://gitlab.com/rouault-team-public/nordlund_et_al.

## Acknowledgements

The authors thank Diane Lalaina for technical assistance; Pierre-Pascal Lenck-Santini, and members of the Epsztein lab for useful discussions; the animal facility headed by Severine Pellegrino, administrative headed by Fanny Pra and inMAGIC platform headed by François Michel platforms of INMED for support. This study was supported by INSERM, by the European Research Council under the European Community’s Seventh Framework Program (ERC-2013-StG-338141_Intraspace to JE), by the ‘Agence National de la Recherche’ (ANRJCJC ANR-17-CE37-0005 to JKG; DEVHIPPO ANR-21-CE16-0005 to JE; ALERT ANR-19-CE37-0016 and MEMNET ANR-22-CE16-0005 to RM) and The Turing Center for Living Systems (CENTURI) under the France 2030 investment plan under the reference ANR-16-CONV-0001 and the Aix-Marseille University Excellence Initiative - A* MIDEX to HR and NL as well as the ‘Institut Universitaire de France’ (junior membership to JKG).

## Authors contribution

Mathilde Nordlund, Conceptualization, Investigation, Software, Formal analysis, Visualization, Writing—original draft, review and editing; Nicolas Levernier & Massimiliano Trippa, Conceptualization, Data curation, Formal analysis, Visualization, Draft review and editing; Romain Bourboulou & Geoffrey Marti, Conceptualization, Data curation, Software, Formal analysis; Rémi Monasson & Hervé Rouault, Conceptualization, Formal analysis, Draft review and editing; Jerome Epsztein, Conceptualization, Resources, Supervision, Funding acquisition, Writing— original draft, review and editing; Julie Koenig-Gambini, Conceptualization, Resources, Supervision, Funding acquisition, Software, Formal analysis, Visualization, Project administration, Writing—original draft, review and editing.

## Supplemental information

**Supplementary Figure 1.**
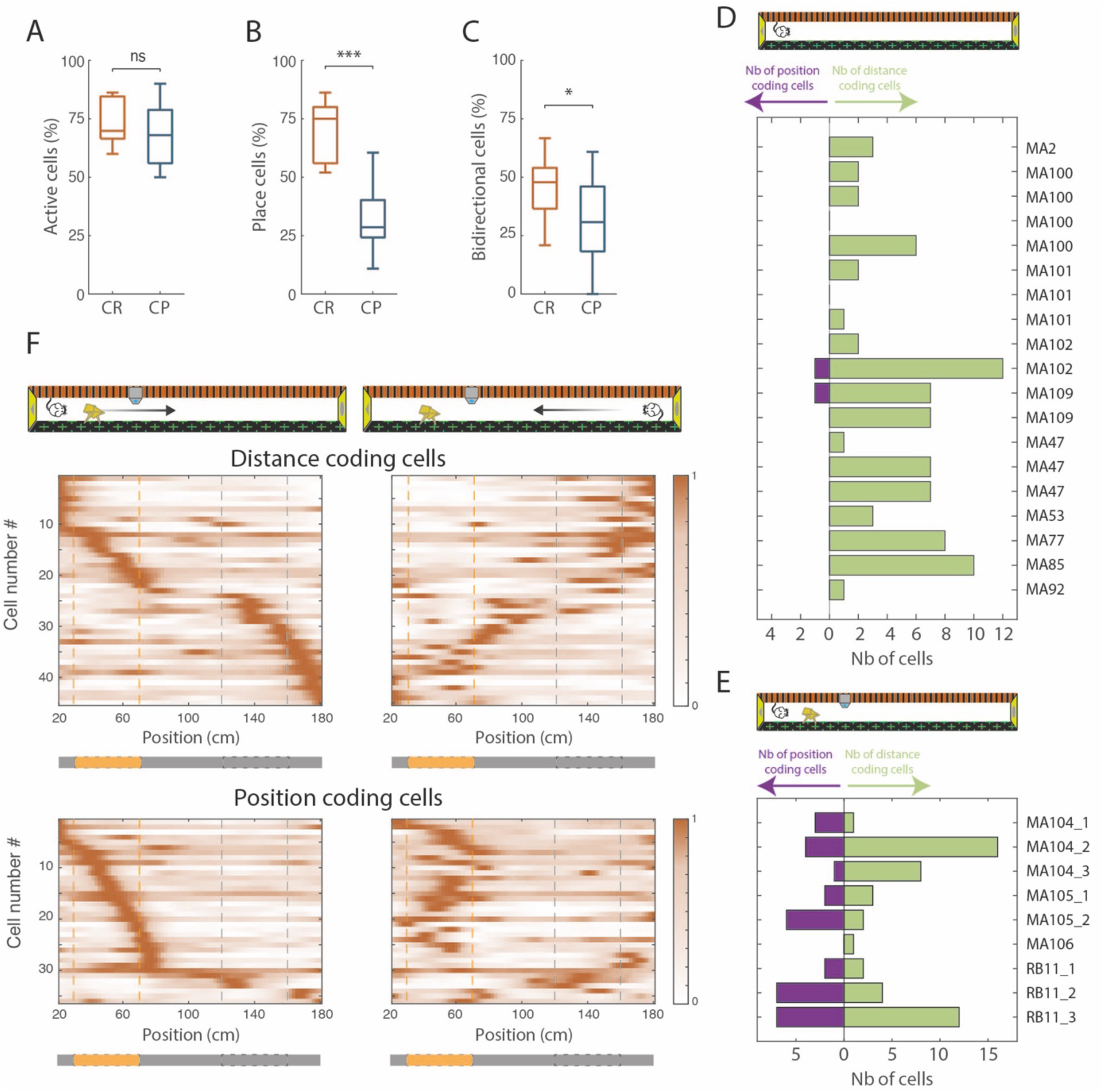
Effects of 3D Objects on the activity and spatial modulation of pyramidal CA1 cells. (**A-C**) Box plots of the percentage of active cells (**A**), place cells (**B**) and bidirectional place cells (**C**) in the cue-rich (CR, red) or cue-poor track (CP, blue). (**D**) Number of position-coding cells (purple) and distance-coding cells (green) for each session recorded in the cue-poor track. (**E**) Number of position-coding cells (purple) and distance-coding cells (green) for each session recorded in the cue-rich track. (**F**) Color-coded mean firing rate maps in the forth (left) and back (right) trials of bidirectional place cells recorded in the cue-rich track and classified as distance-coding cells (top) or position-coding cells (bottom). The color codes for the intensity of the bin’s mean firing rate normalized on the maximal mean firing rate (peak rate) in each direction. The place cells are ordered according to the position of their peak rate in the track for all forth trials (reward zones excluded). Bottom: the track was divided into cue-rich zones (in orange) and cue-poor zones (in gray). The distribution of place fields was compared between a 40 cm cue-rich zone (orange dotted lines) and a 40 cm cue-poor zone (gray dotted lines).

**Supplementary Figure 2:**
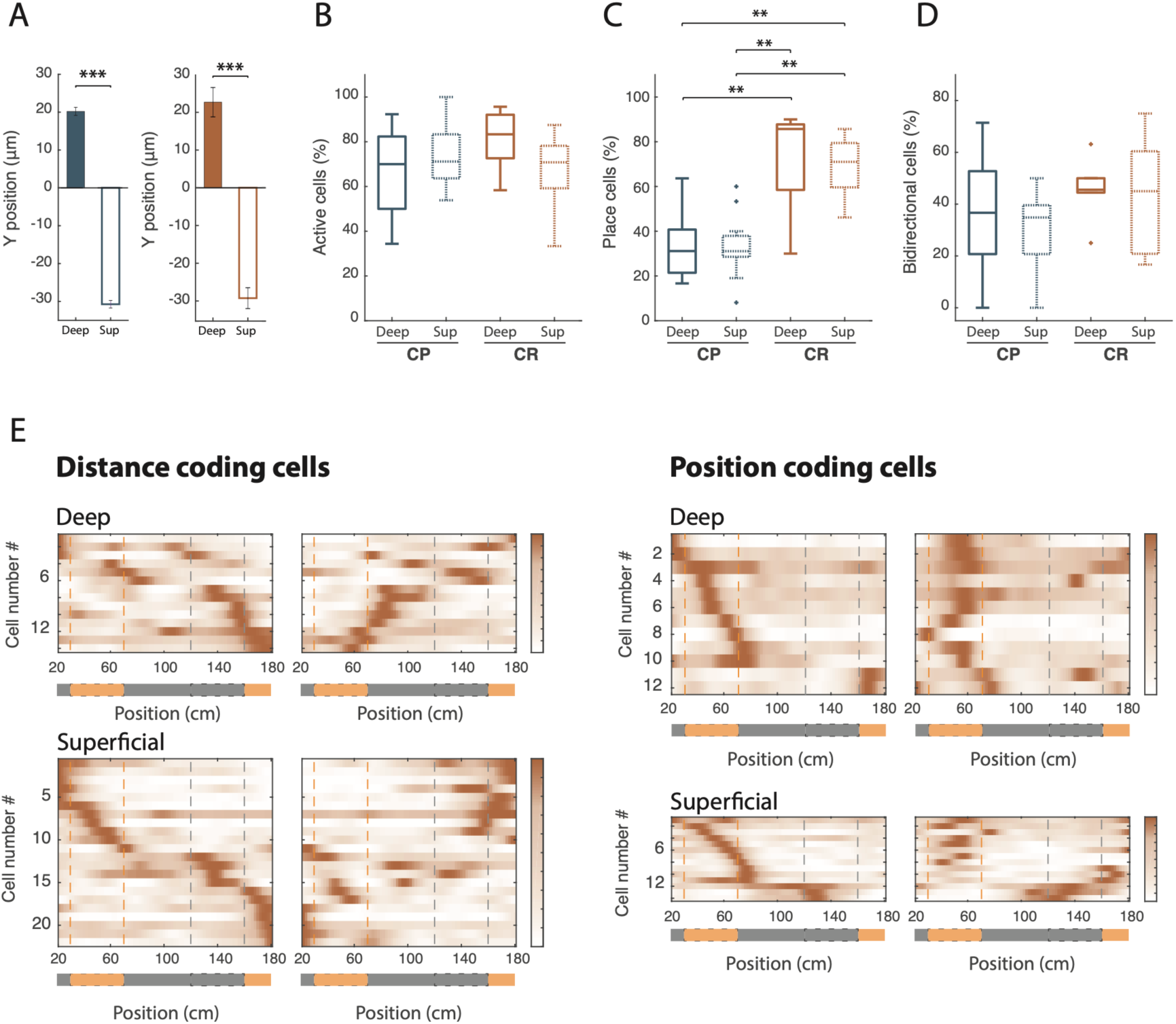
Characterization of deep and superficial firing patterns in the cue-rich and cue-poor tracks. (**A**) Mean ± error type Y position in the CA1 pyramidal cell layer for deep and superficial cells in the cue-poor (blue, left; Z=-26.34, p < 10^-152^, two-tailed WRS test) and cue-rich (red, right; Z=-15.62, p < 10^54^, two-tailed WRS test) tracks. (**B-C**) Box plots of the percentage of deep and superficial active (**B**: X^2^(3) = 3.79, p = 0.29, Kruskal-Wallis test) and place (**C** : X^2^(3) = 22.06, p < 10^4^, Kruskal-Wallis test) cells in the cue-rich (CR, red) and cue-poor track (CP, blue) tracks. (**D**) Box plot of the percentage of deep and superficial bidirectional place cells in the cue-rich (CR, red) and cue-poor track (CP, blue) track (No effect of cell types : F(1,30) = 0.53, p = 0.47; No effect of track condition : F(1,30) = 2.91, p = 0.098 and no interaction between these two factors, F(1,30) = 0.12, p = 0.73, 2-way ANOVA test).(**E**) Color-coded mean firing rate maps in the back-and-forth trials of distance-coding cells (left) for deep and superficial cells and of position-coding cells (right) for deep and superficial cells recorded in the cue-rich track. The color codes for the intensity of the bin’s mean firing rate normalized on the maximal mean firing rate (peak rate) in each direction. Place cells are ordered according to the position of their peak rate in the track for all forth trials (reward zones excluded). Bottom: The track was divided into cue-rich zones (in orange) and cue-poor zones (in gray). The distribution of place fields was compared between a 40 cm cue-rich zone (orange dotted lines) and a 40 cm cue-poor zone (gray dotted lines).

**Supplementary Figure 3:**
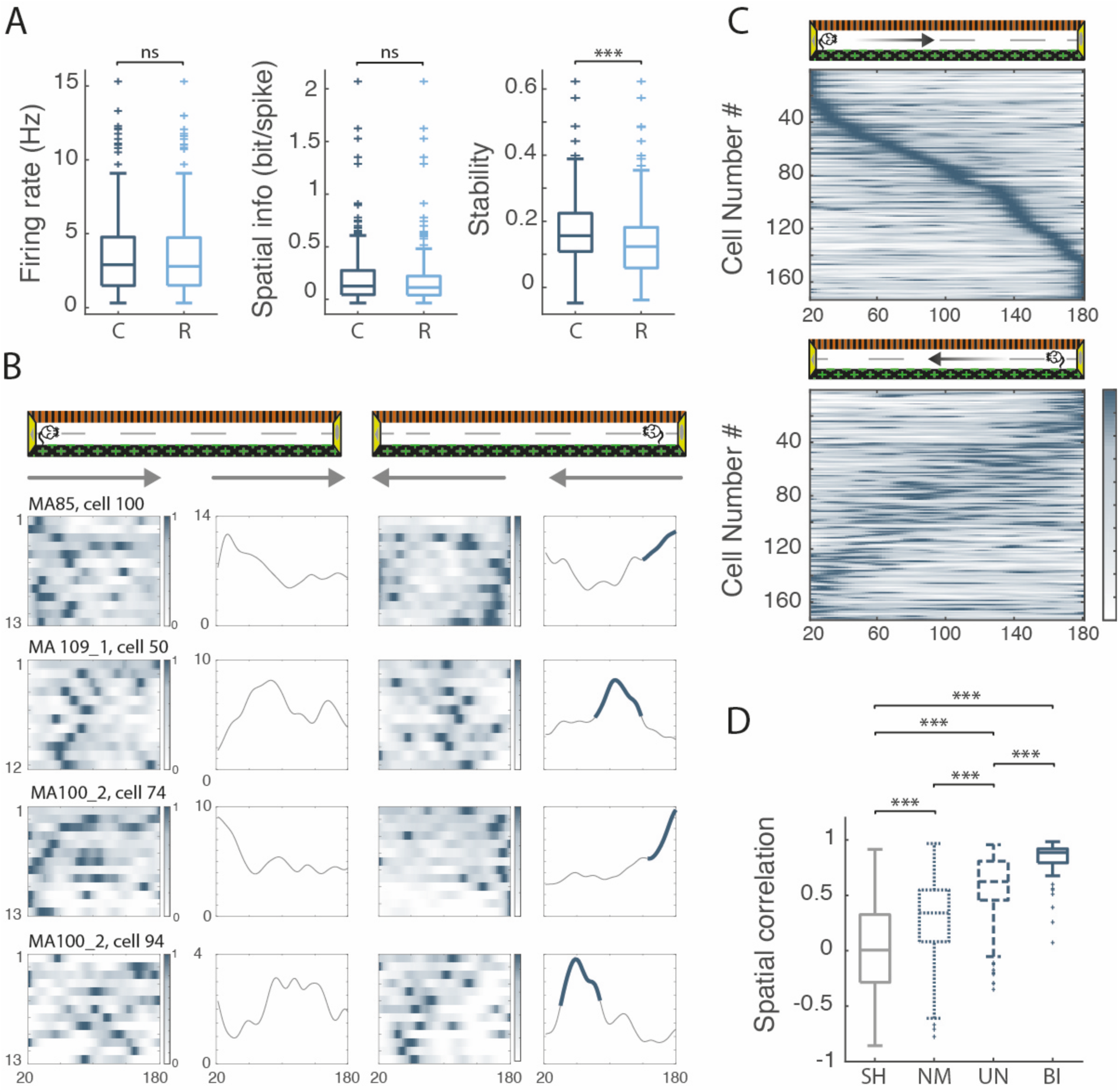
Distance coding in unidirectional place cells. (**A**) Box plots of firing rate (left), spatial information (middle) and stability (right) for both directions: the coding direction (C) and the reverse direction (R). (**B**) 4 examples of unidirectional cells recorded in the cue-poor track. For each cell, the panel shows the color-coded firing rate map for successive trials in both directions (arrows) as a function of the position in the maze and the corresponding mean firing rate by position (reward zones excluded). Bold traces indicate positions of the detected place field. (**C**) Color-coded mean firing rate maps in the forth (top) and back (bottom) trials of all unidirectional cells recorded in the cue-poor track. The color codes for the intensity of the bin’s mean firing rate normalized to the maximum mean firing rate (peak rate) in each direction. The place cells are ordered according to the position of their peak rate in the track for all forth trials (reward zones excluded). Note that the mean firing map in one direction is the mirror image of the mean firing map in the other direction, indicating strong distance coding in unidirectional place cells. (**D**) Box plots of spatial correlation for shuffled pairs (SH = shuffling), non-place cells (NP), unidirectional cells (UN) and bidirectional cells (BI).

**Supplementary Figure 4:**
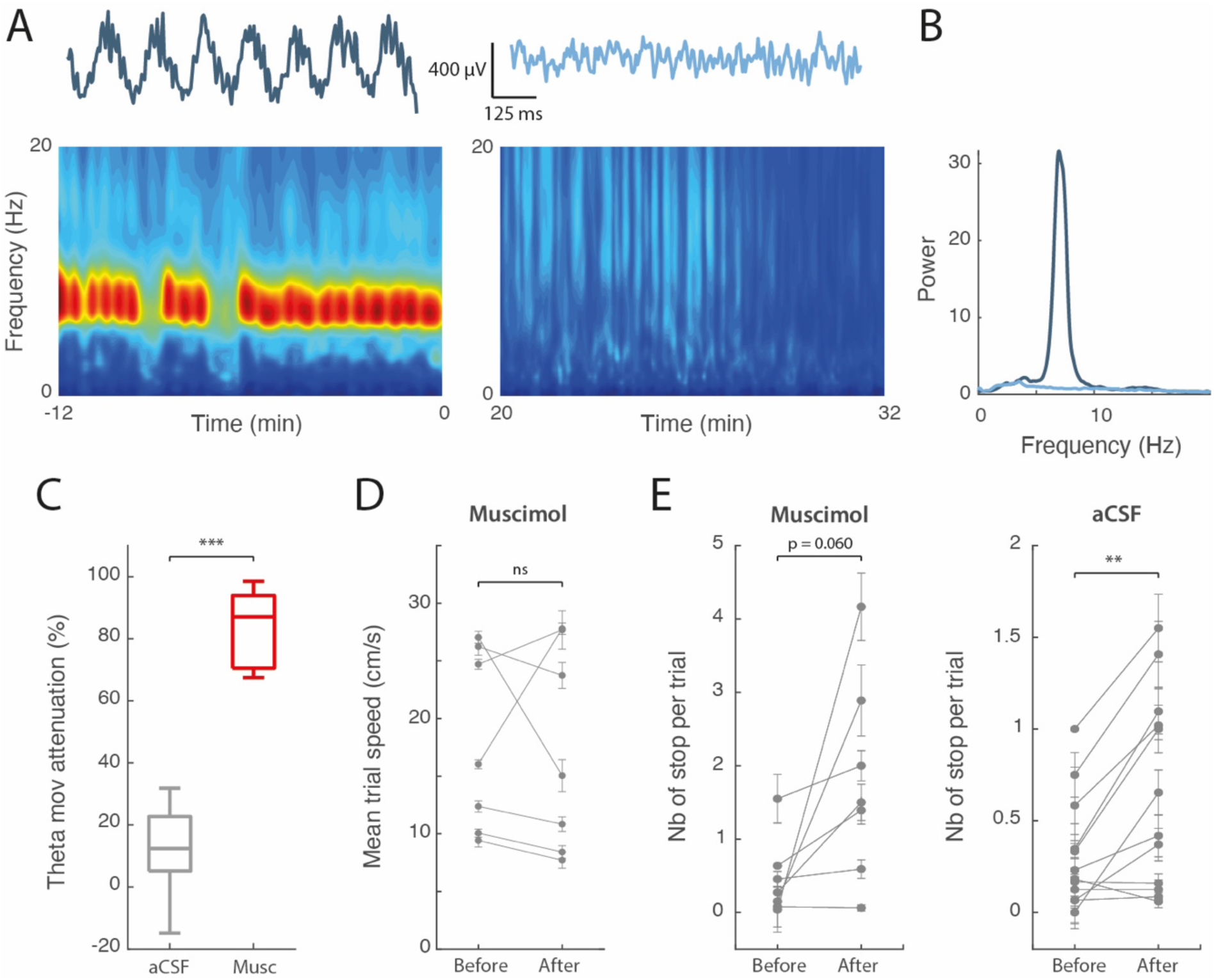
Medial septum inactivation decreased hippocampal theta LFP. (**A**) Spectrograms before (left) and after (right) muscimol infusion into the medial septum (MS) for one recording session. At the top of each spectrogram, 1 second of the raw LFP traces are shown. (**B**) Corresponding power spectrum (left) between 0 and 20 Hz before (dark blue) and during (light blue) MS inactivation. Theta power was reduced by 97.1% during MS inactivation (**C**) Box plots of theta power attenuation during running periods in the control (aCSF) and muscimol groups. (**D**) Mean trial speed (±SEM) before and after muscimol infusion in MS for the 8 recording sessions. (**E**) Mean number of stops per trial before and after the infusion for the muscimol group (left) or the aCSF group (right).

**Supplementary Figure 5:**
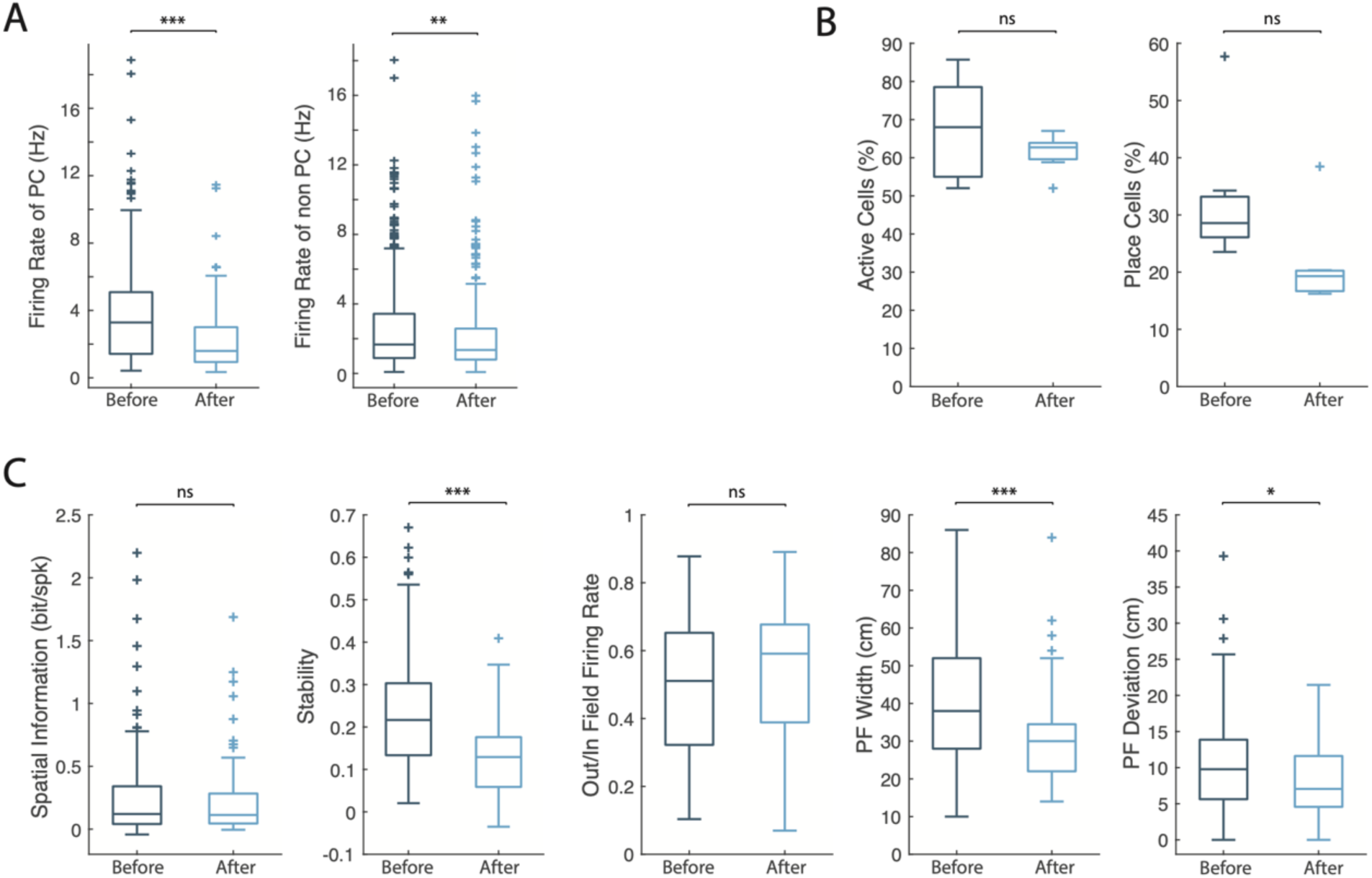
Medial septum inactivation moderately affects hippocampal place cells. (**A**) Box plots of the firing rate of non-place cells (left) or place cells (right) before and after muscimol infusion in the cue-poor track. (**B**) Box plots of percentage of active cells (left) or place cells (right) before and after muscimol inactivation in the cue-poor track. (**C**) Box plots of spatial information, stability, out/in field firing rate, place field width and place field deviation before and during MS inactivation.

**Supplementary Figure 6:**
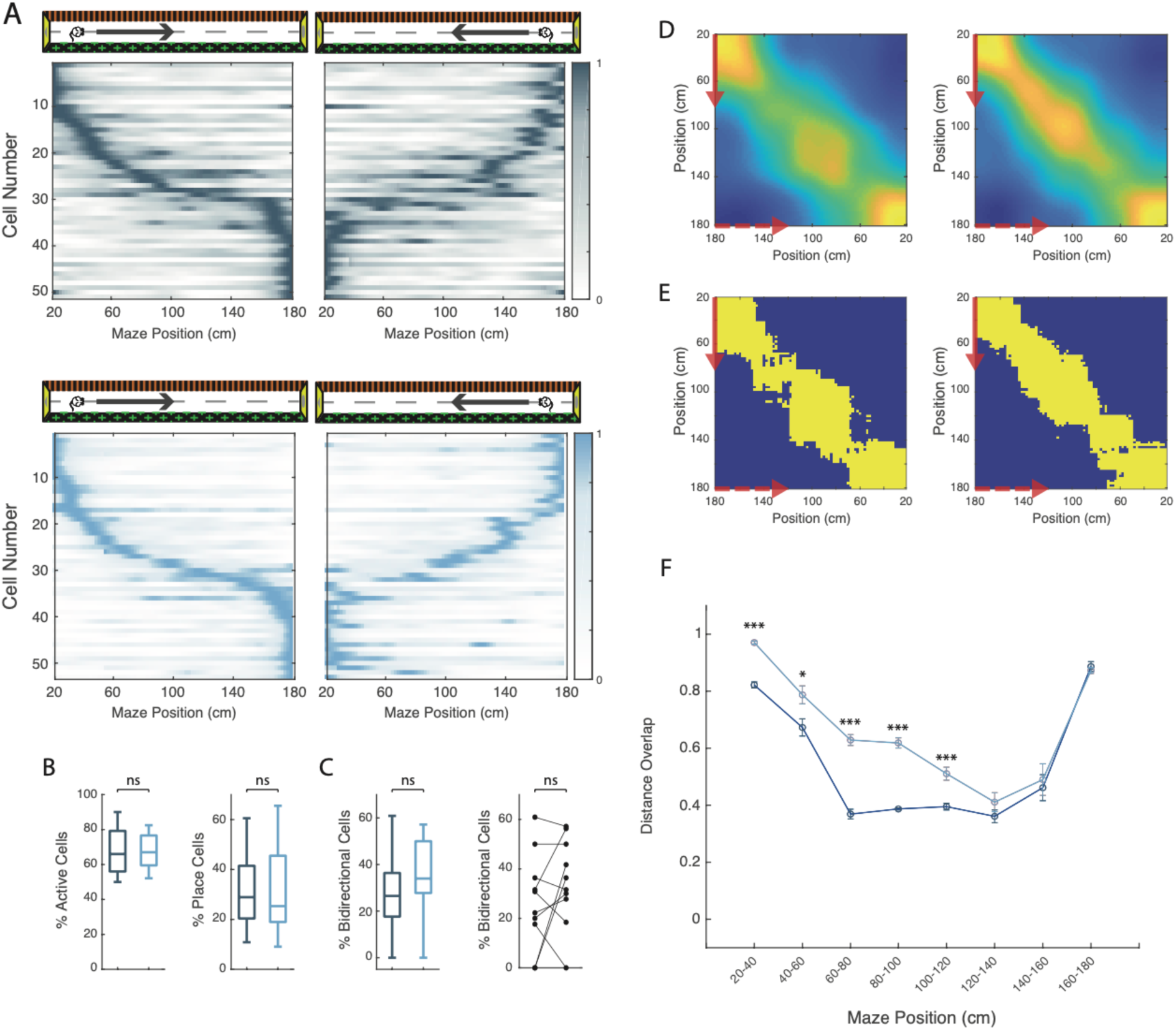
No effect of aCSF injection in the MS on distance coding in bidirectional cells. (**A**) Color-coded mean firing rate maps in the forth (left) and back (right) trials of bidirectional cells recorded before (top, dark blue) and after (bottom, light blue) the aCSF infusion in the cue-poor track. The color codes for the intensity of the bin’s mean firing rate normalized to the maximum mean firing rate (peak rate) in each direction. Pllace cells in both directions are ordered according to the position of their peak rate in the forth trials (reward zones excluded). (**B**) Box plots of the percentage of active cells (left) and place cells (right) before and after aCSF infusion in the medial septum. (**C**) Left: box plots of the percentage of bidirectional cells before and after aCSF infusion in the medial septum; right : percentage of bidirectional cells for each recording session before and after aCSF infusion (**D**) Population vector correlations, at each position’s bin, between place field presence in bidirectional cells recorded in cue-poor track (reward zones excluded) in both directions before (left) and after (right) aCSF infusion. For each axis, the arrows indicate the trial direction (solid red for the forth trials and dashed red for the back trials). The intensity of correlation is color-coded, from the highest (red) to the lowest (blue). (**E**) Population vector matrices showing bins with correlation above the 99^th^ percentile threshold of the shuffled data before (left) and after (right) aCSF infusion in the medial septum. (**G**). Distance overlap of bidirectional place cells in 8 portions of the track in the cue-poor track before (dark blue) and after (light blue) aCSF infusion.

**Supplementary Figure 7:**
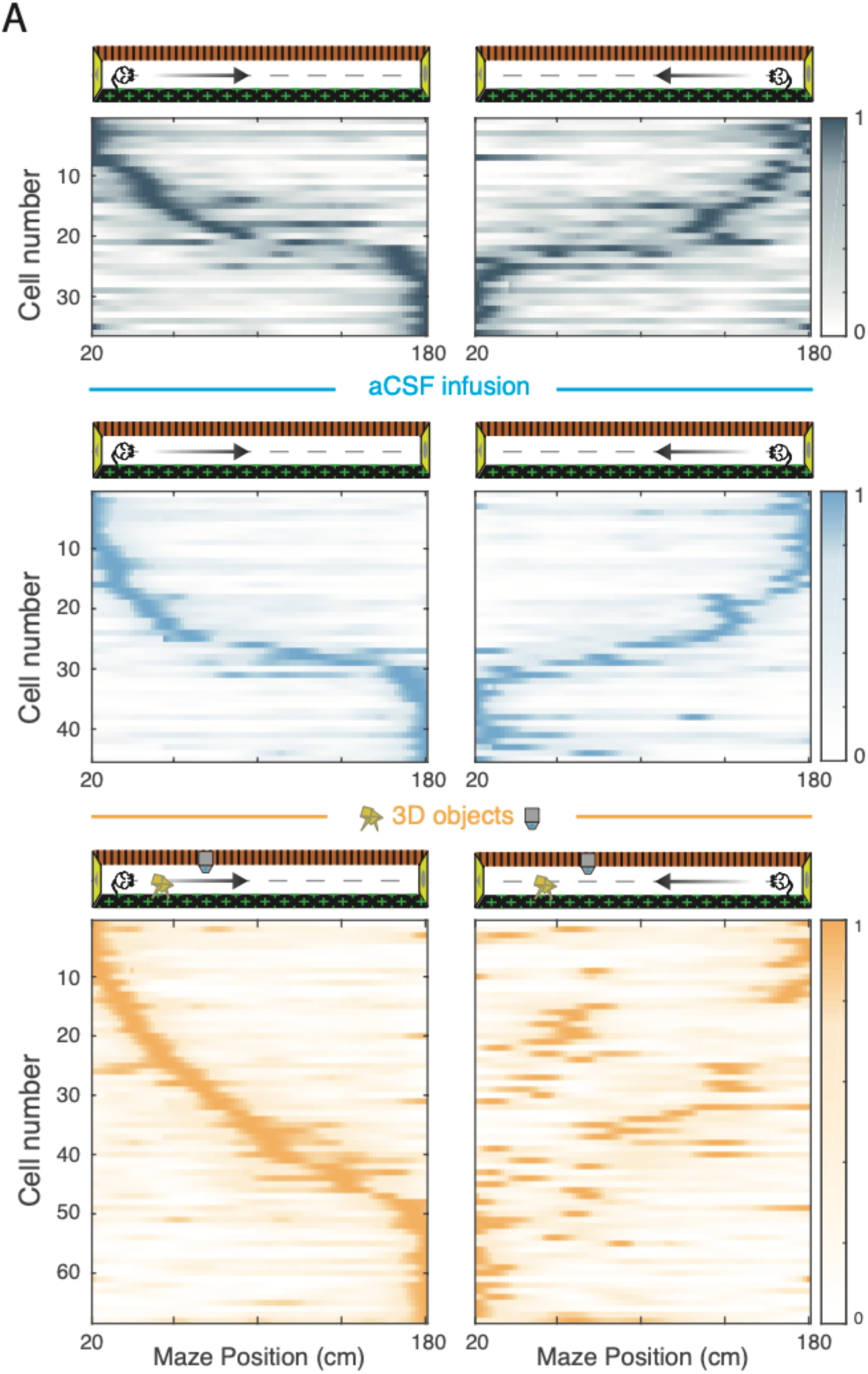
Position and distance coding in the cue-rich track after aCSF infusion. (**A**) Color-coded mean firing rate maps in the forth (left) and back (right) trials of bidirectional cells recorded before (top) and after the muscimol infusion in the cue-poor track (middle) and in the cue-rich track (bottom). The color codes for the intensity of the bin’s mean firing rate normalized to the maximum mean firing rate (peak rate) in each direction. Place cells are ordered according to the position of their peak rate in the track for all forth trials (reward zones excluded).

